# Modulation of Nutritional Composition and Aroma Volatiles in Cultivated Pork Fat by Culture Media Supplementation

**DOI:** 10.1101/2025.07.29.667495

**Authors:** Natsu Sugama, Emily T. Lew, Camilo Riquelme-Guzmán, Di Sheng Lee, Xinxin Li, John S.K. Yuen, Taehwan Lim, Anson Kwan, Run Yi Liu, Yoshene A. Ma, Scott C. Frost, David L. Kaplan

**Affiliations:** Tufts University School of Engineering, Medford, MA 02155, USA; Tufts University School of Arts and Sciences, Medford, MA 02155, USA; Tufts University Cellular Agriculture Commercialization Laboratory, Boston, MA 02111, USA; Deco Labs, Inc, Boston, MA 02111, USA

**Author notes:** **Corresponding Author:** David L. Kaplan.

**Keywords:** Cellular Agriculture, Cultivated meat, Fat, Adipogenesis, Flavor, Aroma, Sensory

## Abstract

Cultivated meat is emerging as a novel food source with the potential to contribute to a more sustainable and ethical food production system. However, limited research to date has explored the extent to which the nutrition and the aroma of such foods can be altered through cell culture conditions. Here, we aimed to modulate the aromatic volatile compounds in heated porcine cultivated fat cells by manipulating the media components while ensuring the preservation of robust fat differentiation. Using dynamic headspace gas chromatography-mass spectrometry (DHS-GC-MS), we demonstrated that supplementing cells with thiamine-HCl increased its intracellular concentration and promoted the production of 4-methyl-5-thiazoleethanol, contributing to milky aroma. Similarly, supplementation with L-methionine enhanced its intracellular concentration and increased the production of methional, a volatile compound with a potato-like aroma. Additionally, myoglobin significantly altered the volatile organic compound profile of cultivated fat. Notably, the concentration of γ-nonalactone, (E,E)-2,4-decadienal and 2-pentylfuran were increased, which contribute to a coconut-like, deep fat, fruity aroma, respectively, as well as elevated levels of other alcohols, aldehydes and furans. These findings highlight the potential of culture media formulations to modulate the aroma in cultivated fat production, a unique opportunity to optimize sensory features using this novel food production technology.

**Highlights:** Nutrient composition and aroma profiles of cultivated pork fat upon baking were modulated by cell culture media supplementation. Supplementing with thiamine-HCl, L-methionine, or myoglobin increased intracellular levels of thiamine or methionine and modulated the formation of aroma volatiles, enhancing characteristic odors such as milky, potato-like, and coconut-like notes.

**Graphical Abstract:** 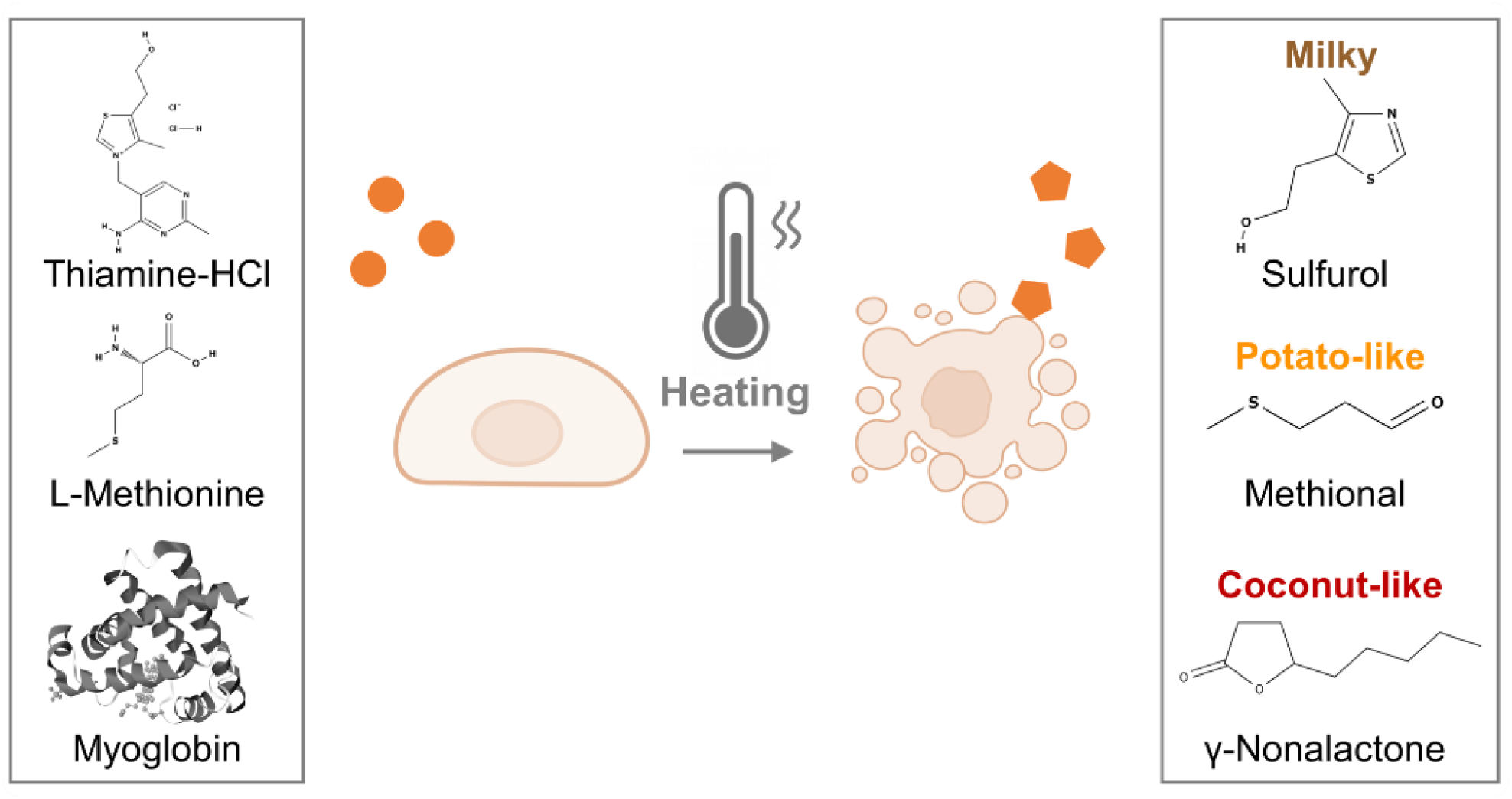

Graphical overview of the methodology. Porcine dedifferentiated fat cells (pDFAT) were differentiated into adipocytes using adipogenesis media supplemented with aroma precursors. The cells were heated (cooked) and the resulting volatile compounds were analyzed using dynamic headspace gas chromatography-mass spectrometry (DHS-GC-MS).

## 1. Introduction

Cultivated meat has emerged as a promising technology to produce meat sustainably, with a significantly reduced risk of infectious diseases along with potentially improved nutrition (1,2). A cornerstone for the success of this technology is flavor, which is a significant factor influencing consumer acceptability and purchasing decisions for meat products (3–6). In terms of flavor, fat plays a crucial role in retaining aroma compounds and contributing to the persistence of scent (7,8).

Additionally, lipid oxidation products, in combination with Maillard compounds, produce a wide variety of aroma compounds in cooked meat (9–12).

Recently, several studies have reported on the aroma profiles of cultivated fat and muscle (13–15). For example, we have characterized volatile organic compounds (VOCs) released during the cooking of cultivated fat derived from porcine dedifferentiated fat cells (pDFAT) (13). Our study revealed the presence of fatty aldehydes such as pentanal, hexanal, octanal, and nonanal which contributed to fatty and buttery aromas. Sensory evaluation showed no statistical difference in response to the cooked cultivated pork fat when compared to traditional livestock-derived pork fat. Another study analyzed the VOCs of porcine adipocytes derived from porcine adipose-derived stem cells (ADSCs) cultured with scaffolds and found that both conventional and cultivated fat shared multiple common VOCs (14). A separate study revealed that porcine fibroblasts and myoblasts cultured in 10% and 15% serum-containing media exhibited significantly higher concentration of thiophenes which impart a meaty aroma than those cells maintained with 1% serum-containing media (15). These results suggested that cultivated meat and fat contain aroma volatiles similar to those found in livestock grown meat and fat, although some differences are present. Additionally, the media composition may influence the types and quantities of VOC profiles produced by baked cells.

In meat science, changing the composition of animal feed can modify VOC profiles, leading to changes in flavor (16–18). Diets with low protein and well-balanced essential amino acids significantly increased the level of 2-heptanone, which has a fruity smell, and 2,3-octanedione, which imparts the characteristic aroma of pork (18). Additionally, post-harvest treatment of cooked ham with thiamine can increase the concentration of 2-methyl-3-furanthiol and bis(2-methyl-3-furyl)-disulfide, which showed a significant difference in taste tests. These results demonstrated that the flavor of meat can be modified by adding certain nutrients to the diet of livestock. It is also possible to change the aroma during secondary processing after harvesting from the animal for processed meats like ham. However, to provoke these types of aroma changes in meat, 60 to 70 days of feeding is required (18,19). Furthermore, the addition of aroma post-harvest requires secondary processes such as curing, and the presence of nitrites or nitrates used in processed meats can impact the changes in aroma (20). An important potential advantage of cultivated meat technology is in the ability to tailor aroma and nutritional content during cell cultivation due to the direct access of the media to the cells (21). Despite this potential, no research to date has specifically targeted the regulation of volatile aroma compounds in cultivated meat or fat through optimization of media composition.

The pathways for VOC generation in meat can be classified into the Maillard reaction, thiamine degradation, lipid oxidation, and Maillard-lipid interactions (22,23). To address these pathways, three additives were studied: (1) thiamine plays a critical role as a coenzyme in carbohydrate metabolism and neural function, and its deficiency is associated with neurodegenerative disorders (24). In addition, during thermal degradation, thiamine generates aroma compounds such as furanthiols (25), thiophenes (26), and thiazoles (27), which contribute to meaty and nutty aromas characteristic of baked meat; (2) L-methionine is an essential amino acid involved in methylation reactions, antioxidant defense via glutathione synthesis, and hepatic function. Upon heating, it undergoes Strecker degradation to produce methional, a well-known aroma compound with a savory, potato-like odor that is commonly found in pork, beef, and chicken (28–32). (3) Myoglobin, which contributes to the generation of aroma compounds, as its iron content catalyzes lipid oxidation reactions (33). Additionally, its presence in plant-based meat increases the formation of lipid oxidation products during heating (34).

Here, we aimed to modulate the volatile compound profile of cultivated pork fat cells by manipulating media components. This approach offers a promising opportunity to leverage media formulation as a tool to enhance the sensory qualities of cultivated fat, thereby advancing applicability in food systems.

## 2. Results

### 2.1. Characterization of Porcine Dedifferentiated Fat (pDFAT) Cells and Optimization of Growth and Adipogenesis Media

To achieve rapid cell proliferation, as well as the maintenance of adipogenic capability over multiple passages and efficient adipocyte differentiation, the media composition for proliferation and adipogenesis were optimized. Cells were cultured using three different growth media formulations which developed based on previous publications (35–38), here in after ‘20%FBS’, ‘20%FBS+ACY’ and ‘15%FBS+bFGF’, in (**Table S1**). The results showed that cells cultured in 20%FBS experienced slower proliferation, with 72.7 hours doubling time at passage 19, leading to the termination of this condition. In contrast, cells cultured with 20%FBS+ACY exhibited an average doubling time of 33.4 hours, while those cultured with bFGF displayed the fastest growth, with an average doubling time of 22.8 hours (**Figure 1A**). Cell diameters were also monitored across passages (**Figure 1B**). Cells cultured in medium containing 20%FBS reached an average diameter exceeding 20.0 μm (the maximum quantification limit), while those cultured in 20%FBS+ACY and 15%FBS+bFGF media maintained smaller diameters, with averages of 14.6 μm and 13.6 μm, respectively.

**Figure 1.**
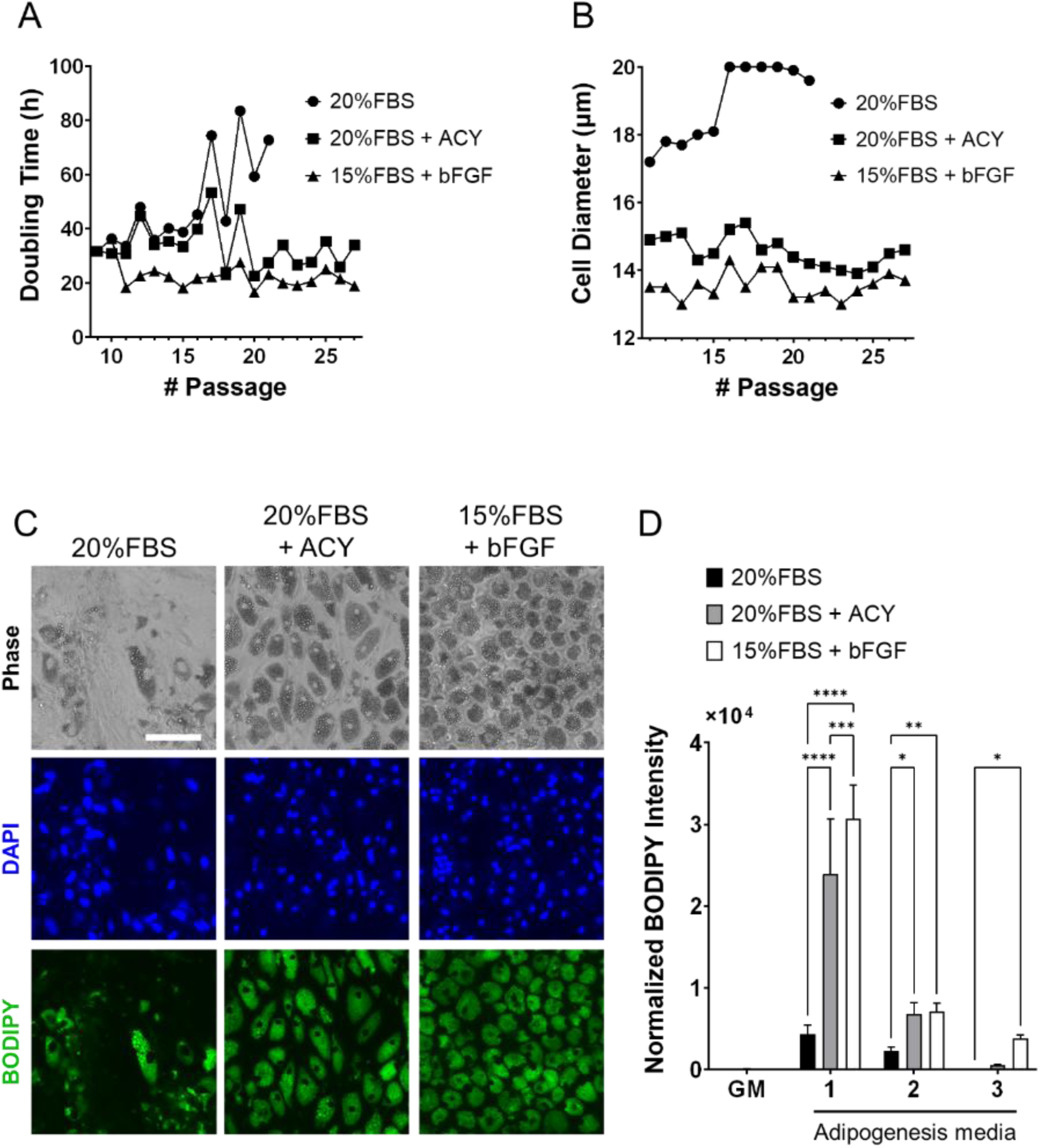
Characterization of pDFAT cells maintained under three different proliferation media, ‘20%FBS’, ‘20%FBS+ACY’ and ‘15%FBS+bFGF’, as well as three different adipogenesis media (1 to 3) to achieve rapid cell growth and maintaining adipogenic capability during continuous passage. **(A)** Hours per cell doubling. **(B)** Cell diameter. **(C)** Morphology of adipocytes maintained with three different proliferation media. Lipids were stained with BODIPY (p23). Scale bars, 100 μm. **(D)** Lipid quantification was performed using BODIPY staining. Average BODIPY integrated intensity was multiplied by BODIPY count and divided by the number of nuclei. GM refers to growth media. Adipogenesis was induced using three different adipogenesis media (Media 1, 2 and 3). n=5 for each group. Statistical significance was determined using Two-way ANOVA with multiple comparisons (*, **, ***, **** denote P < 0.05, P < 0.01, P < 0.001 and P < 0.0001, respectively).

The optimal composition of adipogenesis media were studied. The three different adipogenesis media formulations were based on published media (39–41) (**Table S1**), here in after referred to as “Media1, 2 and 3”. The morphology of the cells maintained in ‘20%FBS’, ‘20%FBS+ACY’ and ‘15%FBS+bFGF’ proliferation media and induced adipogenesis by Adipogenesis Media1 was stained with BODIPY (**Figure 1C**). Among the condition of proliferation media, ‘20%FBS’, ‘20%FBS+ACY’ or ‘15%FBS+bFGF’, and Adipogenesis Media1, 2 and 3 were tested, the combination of cells maintained in ‘15%FBS+bFGF’ proliferation media and induced to undergo adipogenesis using Media1 exhibited the highest lipid accumulation capacity compared to cells cultured under all other growth media conditions (**Figure 1D**). Therefore, the combination of proliferation media ‘15%FBS+bFGF’ and Adipogenesis Media1 was selected for subsequent experiments.

### 2.2. Analysis of Cell Proliferation and Adipogenic Efficiency with Aroma Precursor Supplementation

To investigate the effects of thiamine-HCl or L-methionine on cell proliferation, relative DNA amount was quantified (**Figure 2A**). Additionally, the effects of thiamine-HCl, L-methionine and myoglobin on lipid accumulation were assessed (**Figure 2B**). During the adipogenesis lipid accumulation period, supplementation with thiamine-HCl and L-methionine did not result in a decrease in lipid quantities. However, during the cell proliferation period, supplementation with L-methionine at concentrations above 2.0 mM reduced cell growth. Therefore, all supplements were added only during the adipogenesis lipid accumulation period since there was no beneficial effect of the supplementation on cell proliferation. Optimal concentrations, 500 μM thiamine-HCl, 5.0 mM L-methionine, and 3.0 mg/mL myoglobin were chosen for the supplementation for the rest of this work.

**Figure 2.**
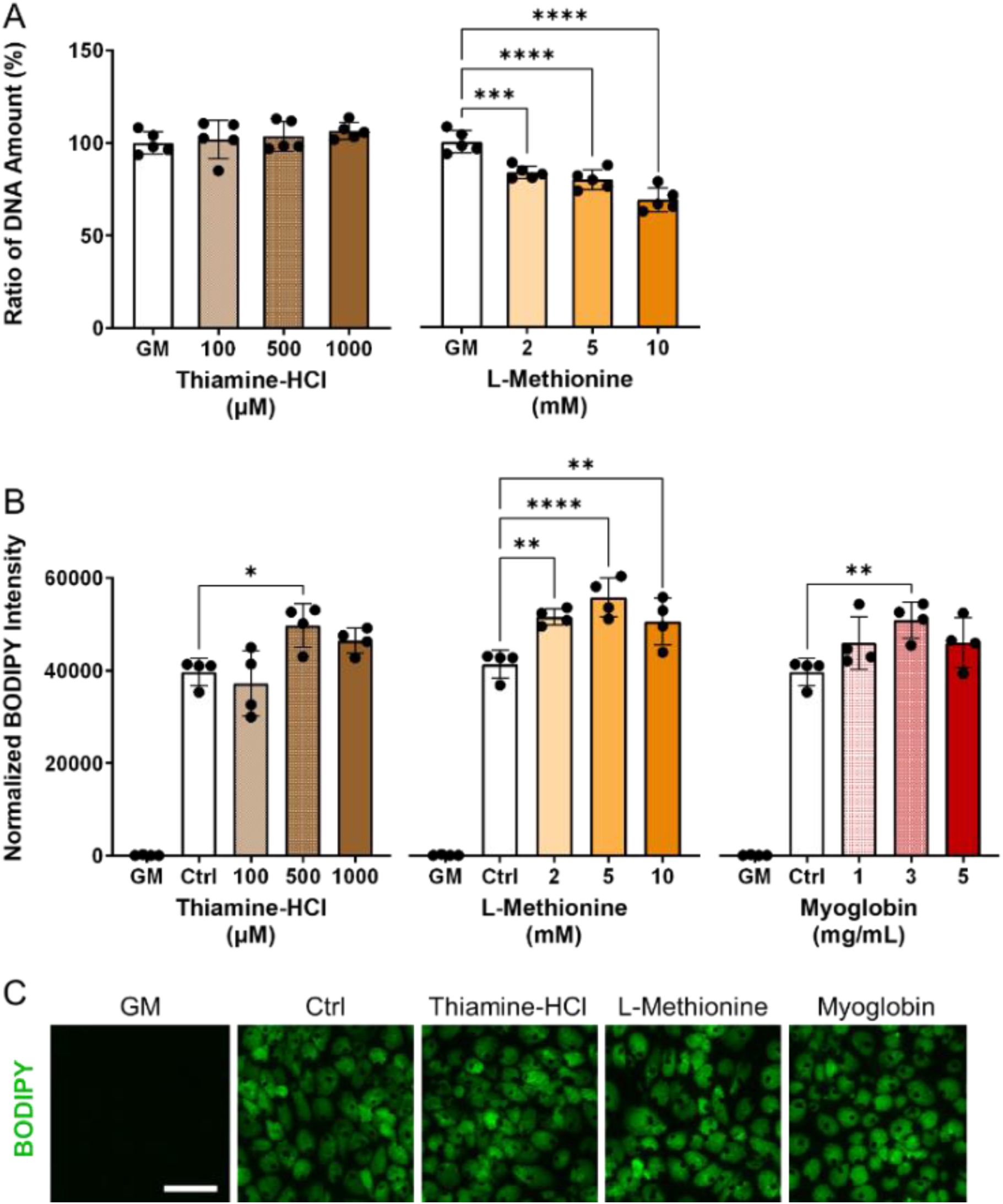
Effect of media supplements on proliferation and adipogenesis in pDFAT cells. **(A)** DNA quantification for pDFAT cells maintained with the proliferation media supplemented with different media additives (n=5). The non-supplemented growth media condition (GM) was considered as 100%. **(B)** Lipid quantification of pDFAT-derived adipocytes treated with supplements during adipogenesis lipid accumulation period. Average BODIPY integrated intensity was multiplied by BODIPY count and divided by the number of nuclei. GM; cultured in proliferation media, Ctrl; cultured in adipogenesis media without supplementation. Statistical significance was determined using one-way ANOVA followed by Tukey’s test (*, **, ***, **** denote P < 0.05, P < 0.01, P < 0.001 and P < 0.0001, respectively) compared to Ctrl. **(C)** Morphology of pDFAT-derived adipocytes cultured in adipogenesis media supplemented with 500 μM Thiamine-HCl, 5 mM L-Methionine or 3 mg/mL Myoglobin. Scale bars, 100 μm.

### 2.3. Metabolite Analysis of Cultivated Fat Before Cooking

To determine whether thiamine-HCl or L-methionine were taken up by the cells, a metabolite analysis was performed. The relative amounts of thiamine, thiamine pyrophosphate, methionine, methionine sulfoxide, S-adenosyl-L-homocysteine (SAH), and S-adenosyl-L-methionine (SAM) were analyzed. An increase in thiamine pyrophosphate, a downstream intermediate of thiamine, was detected (**Figure 3A**). In addition, the levels of SAH, the immediate, and SAM, a subsequent downstream intermediate of L-methionine, were both elevated when compared to the non-supplemented control (**Figure 3B**). Methionine sulfoxide, an oxidative product of L-methionine (42), was increased. The metabolic pathways of thiamine and L-methionine of pig (*Sus scrofa*) were confirmed using the KEGG database (ssc00730, ssc00270)

**Figure 3.**
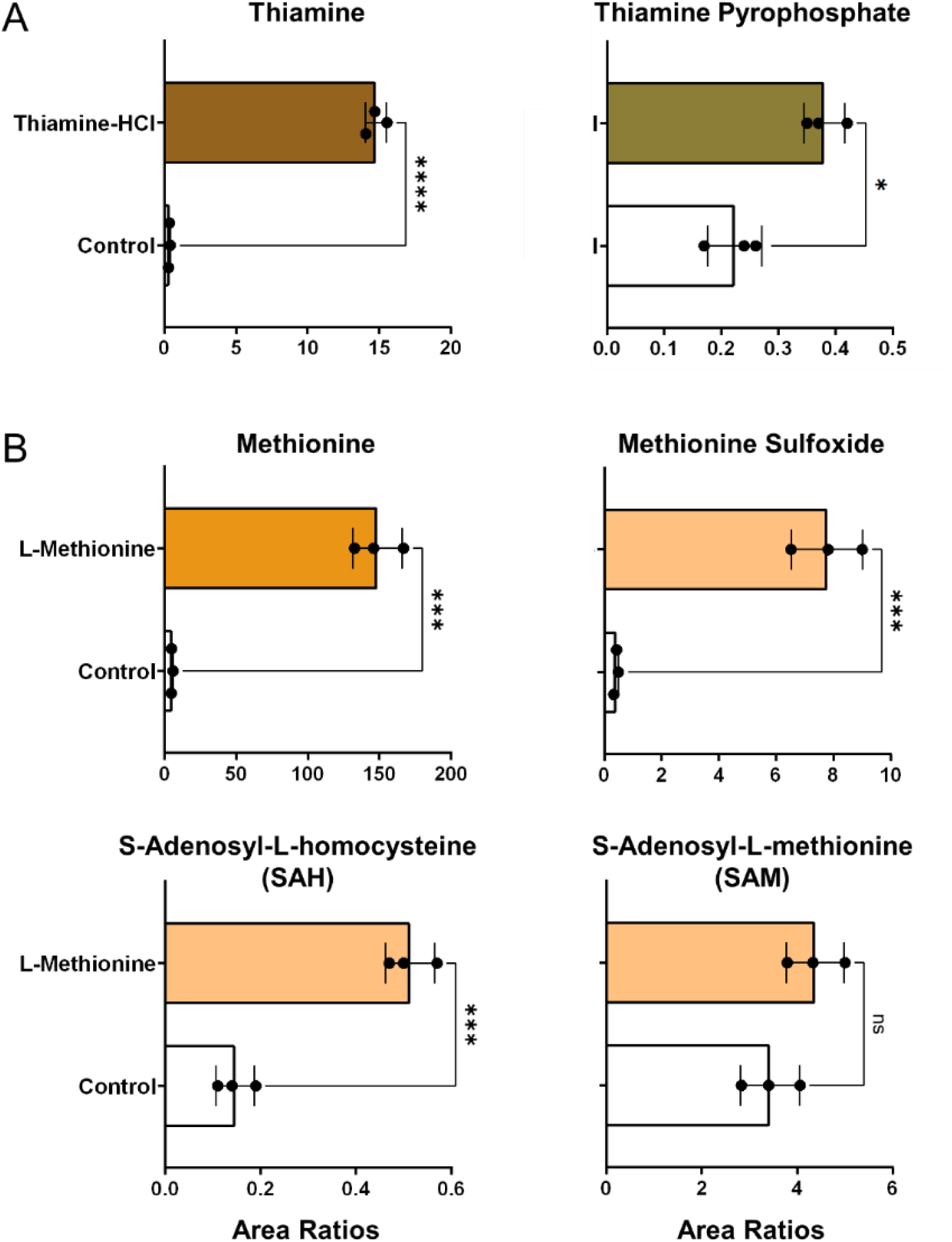
Relative levels of metabolites in cultivated fat. **(A)** Peak area ratios of thiamine and thiamine pyrophosphate in non-supplemented control and thiamine-HCl-supplemented cells prior to baking (mean±SD, n=3). **(B)** Peak area ratios of methionine and its downstream intermediates in non-supplemented control and L-methionine-supplemented samples prior to baking (mean±SD, n=3). Area ratios were calculated using the following formula: [{(Target Peak Area)/(Heavy Carbon-Labeled Phenylalanine or Methionine) divided by protein content measured by BCA assay. Statistical significance was determined by unpaired t-test (*, ***, **** denotes P < 0.05, 0.001, 0.0001, respectively.

### 2.4. Fatty Acid Analysis of Harvested Fat Before Cooking

To determine the specific types of fatty acids accumulated as triglycerides or phospholipids in cultivated fat, fatty acid analysis was performed. Additionally, the fatty acid profiles of both non-supplemented and myoglobin-supplemented cultivated fat were analyzed prior to baking, as the iron in myoglobin could potentially influence the composition. The results showed that the fatty acid composition of the non-treated samples consisted of 45.6% C18:1 (cis-9), 17.1% C18:2 (all-cis-9,12), 14.2% C16:0, and 9.89% C18:0, as the top four fatty acids identified (**Figure 4A**). Myoglobin-treated cultivated fat showed a statistically significant decrease of 4.07% in unsaturated fatty acid content, with a corresponding increase in saturated fatty acids **(Figure 4B**).

**Figure 4.**
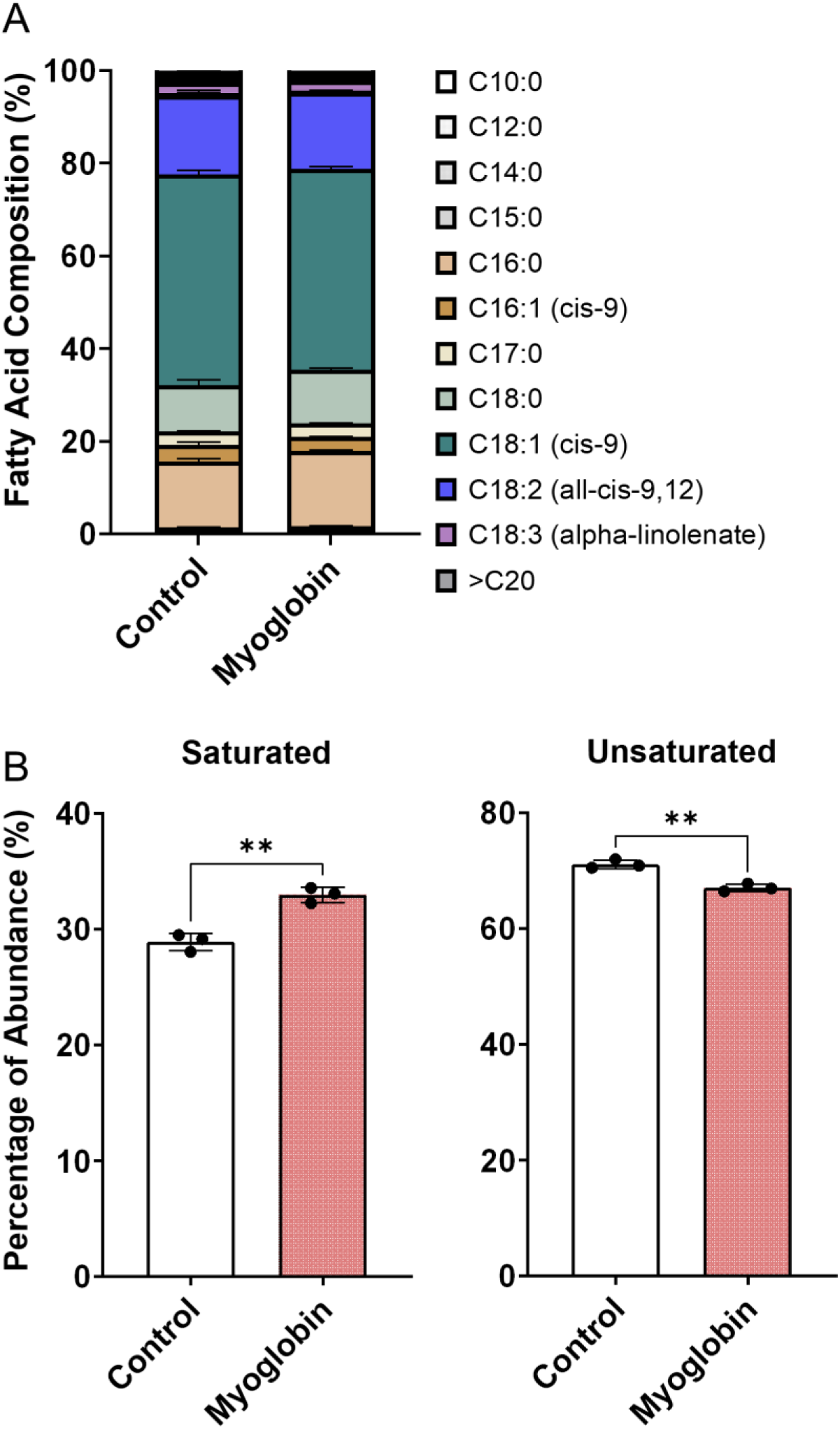
Fatty acid profile of cultivated fat. **(A)** Fatty acid composition between non-treated control and myoglobin, prior to baking (mean±SD, n=3). **(B)** Percentage of total saturated and unsaturated fatty acids. Statistical significance was determined by unpaired t-test (** denotes P < 0.01).

### 2.5. Impact of Aroma Precursor Supplementation on the Concentration and Profiles of Volatile Compounds from Cells

The VOCs produced upon the addition of aroma precursors to the medium and subsequent heating/cooking were analyzed using DHS-GC-MS. The peaks of all compounds identified through deconvolution, were normalized using the internal standards and cell mass, described in ‘GC/MS data processing’ were represented in dot plots. Thiamine-HCl significantly induced the generation of 4-methyl-5-thiazoleethanol (sulfurol), milky aroma compound which derived from thiamine degradation (43,44) (**Figure 5A**). L-methionine promoted the formation of methional, a well-known potato-like aroma compound (**Figure 5B**). The addition of myoglobin caused significant changes to the VOC profile, beginning with the formation of γ-nonalactone, (E,E)-2,4-decadienal, 2-pentylfuran, δ-decalactone, and benzeneacetaldehyde, which are responsible for the coconut-like, deep fat, fruity, peachy, and honey aroma of meat (45–48) , while increasing heptanal and 1-pentanol which imparts green, fatty aroma (49,50) (**Figure 5C**). Furthermore, myoglobin enhanced the production of various lipid degradation products, including aldehydes, alcohols, furans and some of ketones, fatty acids, and hydrocarbons, and phenolic derivatives. Some of the esters also showed an increase; however, the average fold change was not as significant compared to that of aldehydes, alcohols, and furans (**Table 3**). No significant changes were observed in the levels of pyrroles and thiazoles (**Figure 6**). All compounds and peaks found in the non-supplemented controls, along with those supplemented with thiamine-HCl, L-methionine and myoglobin are shown in (**Table 1, 2, 3**).

**Figure 5.**
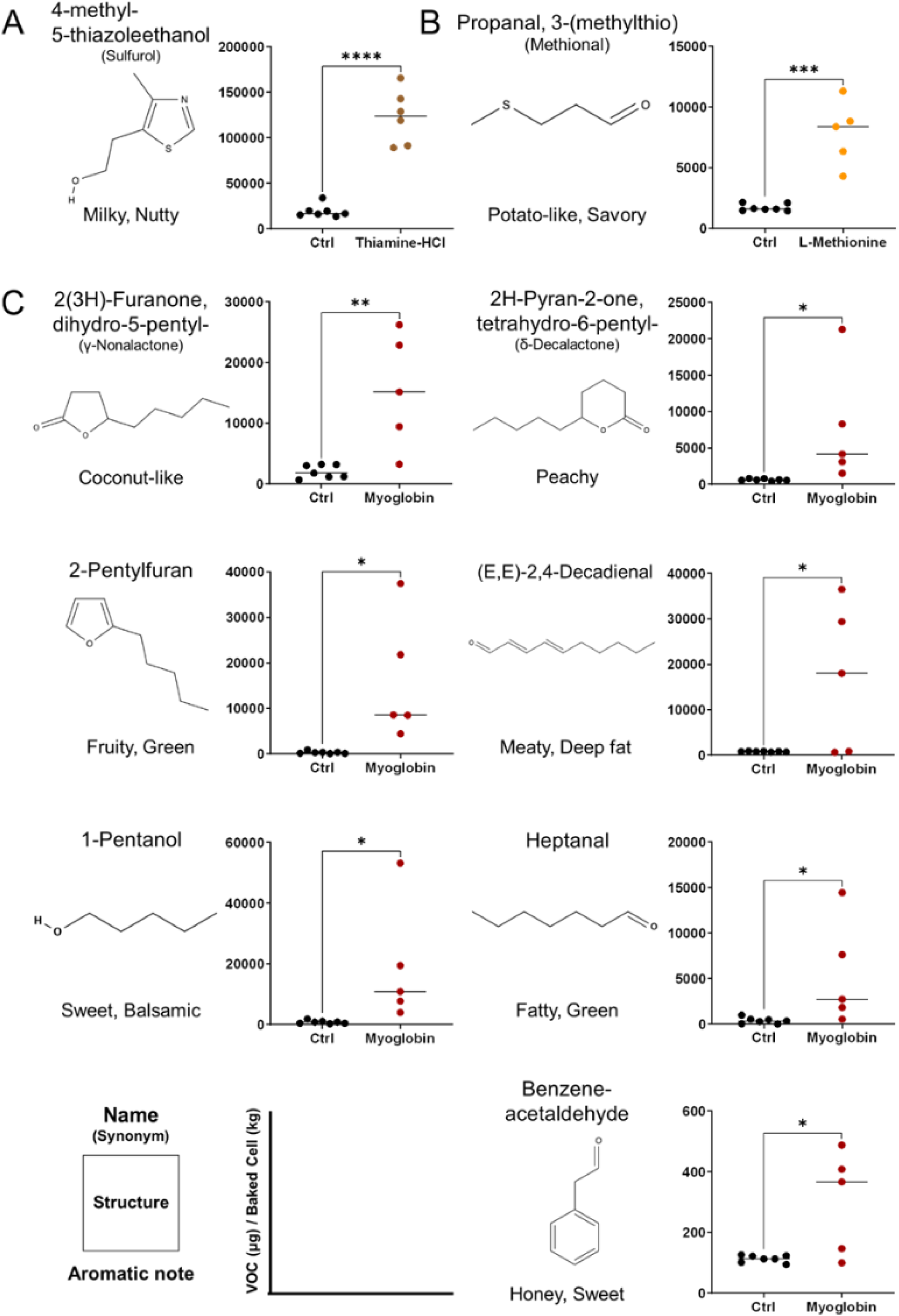
Major volatile organic compounds (VOCs) provoked by media supplementations and detected by DHS-GC-MS derived from pDFAT-derived adipocytes upon baking. VOCs altered under **(A)** 500 μM of Thiamine-HCl, **(B)** 5 mM of L-Methionine, **(C)** 3 mg/mL Myoglobin-supplemented conditions. VOCs were quantified by normalizing peak areas to two internal standards and converting the values to mass using authentic standard curves. The resulting concentrations were further normalized to cell mass after baking. ’Ctrl’ indicates the non-supplemented cell condition. Replicates 5 to 7 include biological triplicates. Statistical significance was determined using unpaired t-test (*, **, ***, **** denote P < 0.05, P < 0.01, P < 0.001 and P < 0.0001, respectively). Compounds were identified by retention index (RI) and Mass Spec referencing NIST17 compared with authentic standards.

**Figure 6.**
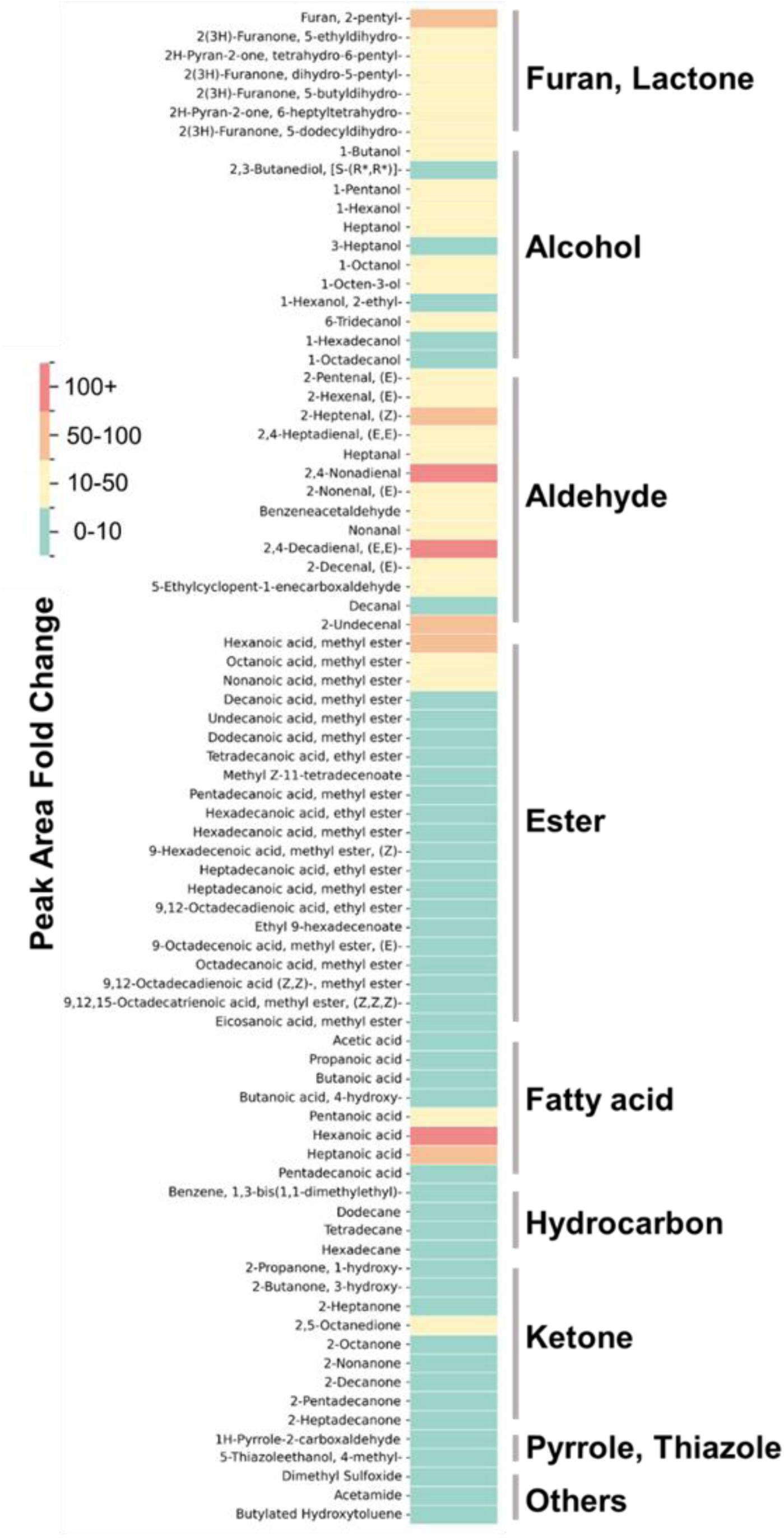
The fold change in the normalized peak area of VOCs found in myoglobin-supplemented cultivated fat, compared to non-supplemented control cells. The color intensity represents the value of the fold changes. All VOCs are tentatively identified except for VOCs shown in Figure3.

**Table 1.** All the Volatile Organic Compounds (VOCs) found in 120℃ heated cultivated fat maintained with non-supplemented media (Control) and supplemented with 500 μM thiamine-HCl. a) Normalized peak area was calculated by the following formula: [{(Peak area of target compound)/(Peak area of 2-Methylheptan-3-one)}*Average peak area of 2-Methylheptan-3-one]/(Peak area of Naphthalene-d8)*(Average peak area of Naphthalene-d8) and divided by the dried cell mass (mg) after baking. b) Statistical significance was determined using unpaired t-test (*, **** denote P < 0.05 and P < 0.0001, respectively). c) All VOCs, except for highlighted in blue, were tentatively identified with both Mass Spec (MS) data and Retention Index (RI). Regarding the compounds shown as ‘MS, RI (standard)’ in the ‘identification’ column, RI was obtained by the authentic standard injection. RI of the other VOCs were taken from the literature which used DB-Wax column, these are considered as tentatively identified.

|  |  | Normalized Peak Area<sup>a</sup><br>(Average ± SD) |  |  |  | RI<sup>c</sup> |  | m/z<br>(actual) |  |  | m/z<br>(NIST) |  |  |  |
| --- | --- | --- | --- | --- | --- | --- | --- | --- | --- | --- | --- | --- | --- | --- |
| Name | CAS# | Control | Thiamine-HCl | p value<sup>b</sup> |  | actual | ref | 1 | 2 | 3 | 1 | 2 | 3 | Identification |
| 4-methyl-5-thiazoleethanol | 504-78-9 | 34801 ± 29213 | 596494 ± 149490 | 0.0000009 | **** | 2268 | 2271 | 112 | 113 | 143 | 112 | 113 | 143 | RI (standard), MS |
| 2-Propanone, 1-hydroxy- | 1638-16-0 | 39789 ± 14488 | 64125 ± 16195 | 0.0155 | * | 1275 | 1278 | 43 | 74 | 42 | 43 | 31 | 74 | RI (literature), MS |
| Dodecanoic acid, methyl ester | 111-82-0 | 11640 ± 17303 | 54750 ± 46633 | 0.0433 | * | 1788 | 1770 | 74 | 87 | 43 | 74 | 87 | 41 | RI (literature), MS |
| 9,12-Octadecadienoic acid, ethyl ester | 7619-08-1 | 7345 ± 7247 | 16766 ± 10711 | 0.0863 | ns | 2506 | 2515 | 81 | 67 | 95 | 67 | 81 | 55 | RI (literature), MS |
| Benzaldehyde | 100-52-7 | 25877 ± 8698 | 37080 ± 12813 | 0.0882 | ns | 1487 | 1502 | 106 | 105 | 77 | 77 | 106 | 105 | RI (literature), MS |
| 9,12,15-Octadecatrienoic acid, methyl ester, (Z,Z,Z)- | 301-00-8 | 63244 ± 72270 | 147952 ± 98005 | 0.1006 | ns | 2537 | 2550 | 79 | 95 | 67 | 79 | 67 | 95 | RI (literature), MS |
| Hexadecanoic acid, methyl ester | 112-39-0 | 824531 ± 864907 | 1662487 ± 890549 | 0.1138 | ns | 2204 | 2207 | 74 | 87 | 143 | 74 | 87 | 43 | RI (literature), MS |
| Ethyl 9-hexadecenoate | 54546-22-4 | 6735 ± 5144 | 13282 ± 8501 | 0.1150 | ns | 2265 | 2267 | 55 | 88 | 69 | 55 | 88 | 69 | RI (literature), MS |
| 9,12-Octadecadienoic acid (Z,Z)-, methyl ester | 112-63-0 | 399566 ± 456548 | 855335 ± 517402 | 0.1194 | ns | 2472 | 2488 | 81 | 67 | 95 | 67 | 81 | 95 | RI (literature), MS |
| 9-Octadecenoic acid, methyl ester, (E)- | 1937-62-8 | 796200 ± 866043 | 1452439 ± 641588 | 0.1548 | ns | 2433 | 2445 | 264 | 265 | 222 | 264 | 97 | 96 | RI (literature), MS |
| Thiazole, tetrahydro- | 13019-20-0 | 3105 ± 5474 | 16242 ± 23483 | 0.1763 | ns | 1454 | 1454 | 89 | 88 | 43 | 89 | 43 | 42 | RI (literature), MS |
| Tetradecane | 629-59-4 | 71370 ± 17261 | 95141 ± 47750 | 0.2429 | ns | 1397 | 1400 | 57 | 71 | 43 | 57 | 43 | 71 | RI (literature), MS |
| 2-Decanone | 513-86-0 | 75638 ± 71964 | 41813 ± 26916 | 0.3022 | ns | 1478 | 1476 | 58 | 43 | 71 | 58 | 43 | 71 | RI (literature), MS |
| Butylated Hydroxytoluene | 128-37-0 | 8811 ± 15390 | 51760 ± 112526 | 0.3359 | ns | 1891 | 1902 | 205 | 220 | 177 | 205 | 220 | 57 | RI (literature), MS |
| 2-Butanone, 3-hydroxy- | 3658-77-3 | 245097 ± 195382 | 342392 ± 173977 | 0.3672 | ns | 1264 | 1265 | 45 | 43 | 88 | 45 | 43 | 88 | RI (literature), MS |
| 2-Nonanone | 705-86-2 | 75255 ± 71218 | 46516 ± 23726 | 0.3676 | ns | 1374 | 1379 | 58 | 43 | 71 | 42 | 58 | 41 | RI (literature), MS |
| 2,3-Butanediol | 104-61-0 | 23896 ± 32533 | 12507 ± 10666 | 0.4317 | ns | 1530 | 1544 | 45 | 57 | 43 | 45 | 43 | 57 | RI (literature), MS |
| Octanal | 124-13-0 | 14679 ± 19686 | 8008 ± 4649 | 0.4372 | ns | 1272 | 1277 | 56 | 57 | 43 | 43 | 44 | 56 | RI (literature), MS |
| 2-Octanone | 821-55-6 | 25557 ± 17099 | 32143 ± 14852 | 0.4780 | ns | 1269 | 1278 | 58 | 43 | 71 | 43 | 58 | 41 | RI (literature), MS |
| 2-Heptanone | 693-54-9 | 67854 ± 44580 | 87608 ± 54498 | 0.4867 | ns | 1173 | 1174 | 43 | 58 | 71 | 43 | 58 | 71 | RI (literature), MS |
| 2(3H)-Furanone, 5-ethylidihydro- | 71-41-0 | 9013 ± 6490 | 6922 ± 3186 | 0.4893 | ns | 1665 | 1665 | 85 | 57 | 56 | 85 | 29 | 56 | RI (literature), MS |
| Furan, 2-pentyl- | 3777-69-3 | 12447 ± 10782 | 9907 ± 7602 | 0.6391 | ns | 1213 | 1215 | 81 | 82 | 138 | 81 | 82 | 132 | RI (literature), MS |
| Dimethyl Sulfoxide | 67-68-5 | 156198 ± 144743 | 193834 ± 166644 | 0.6711 | ns | 1560 | 1553 | 63 | 78 | 61 | 63 | 78 | 45 | RI (literature), MS |
| 4-Octanone | 589-63-9 | 3104 ± 2805 | 3694 ± 2823 | 0.7132 | ns | 1215 | 1231 | 71 | 57 | 85 | 43 | 57 | 71 | RI (literature), MS |
| Tetradecanoic acid, ethyl ester | 124-06-1 | 5232 ± 4731 | 5990 ± 3893 | 0.7609 | ns | 2035 | 2040 | 88 | 101 | 55 | 88 | 101 | 43 | RI (literature), MS |
| Dodecane | 112-40-3 | 7088 ± 11036 | 5478 ± 7491 | 0.7683 | ns | 1192 | 1199 | 57 | 43 | 71 | 57 | 43 | 71 | RI (literature), MS |
| 2-Octenal, (E)- | 111-13-7 | 5506 ± 2251 | 5959 ± 3295 | 0.7747 | ns | 1409 | 1416 | 70 | 55 | 41 | 41 | 55 | 29 | RI (literature), MS |
| Heptanal | 111-71-7 | 11392 ± 4561 | 10958 ± 4406 | 0.8655 | ns | 1174 | 1171 | 70 | 44 | 55 | 70 | 41 | 44 | RI (literature), MS |
| 1-Pentanol | 71-41-0 | 11649 ± 6937 | 12243 ± 8824 | 0.8943 | ns | 1243 | 1255 | 42 | 55 | 70 | 42 | 55 | 41 | RI (literature), MS |
| 2-Pentadecanone | 2548-87-0 | 16462 ± 3256 | 15945 ± 11927 | 0.9138 | ns | 2002 | 2002 | 58 | 43 | 59 | 58 | 43 | 59 | RI (literature), MS |
| Hexadecane | 544-76-3 | 25382 ± 15849 | 24715 ± 6722 | 0.9256 | ns | 1596 | 1600 | 57 | 71 | 85 | 57 | 43 | 71 | RI (literature), MS |
| Nonanal | 124-19-6 | 32805 ± 13222 | 32214 ± 13871 | 0.9388 | ns | 1377 | 1379 | 57 | 56 | 43 | 57 | 41 | 43 | RI (literature), MS |
| Pentadecane | 629-62-9 | 12416 ± 8583 | 12505 ± 4808 | 0.9826 | ns | 1497 | 1500 | 57 | 71 | 43 | 57 | 43 | 71 | RI (literature), MS |

## 3. Discussion

In this study, we demonstrated that aroma volatiles in cultivated fat cells can be altered by media supplementation, resulting in different flavors in fat, while maintaining rapid proliferation and robust adipogenesis capability. Prior to the investigation of flavor precursor supplementation, we aimed to select optimal media to achieve rapid cell proliferation between the three proliferation media previously reported (37,38,51). ‘20% FBS’ was originally used when pDFAT cells were first reported (38) and media supplemented with bFGF has been used in other pDFAT studies and showed robust adipogenic support until around passage 50 (38). While bFGF has a positive effect on both proliferation and differentiation for the cells (52,53), the cost has been highlighted as an issue (54).

Therefore, as an alternative we considered the addition of small molecule compounds, as previously reported in studies with MSCs (37). An improvement in cell proliferation rate was observed with the addition of bFGF or ACY, which aligns with previous reports on pDFAT or human MSCs (**Figure 1A**) (37,38). Additionally, cells maintained in ‘20%FBS+ACY’ or ‘15%FBS+bFGF’ exhibited smaller cell sizes compared to those maintained in ‘20%FBS’ (**Figure 1B**). Several prior studies have reported that aged cells tend to have larger sizes, whereas stem cells or those undergoing rapid self-renewal cycles are typically smaller (55,56), consistent with the results reported here, where the cells maintained with ‘20%FBS+ACY’ or ‘15%FBS+bFGF’ showed faster proliferation and higher adipogenic capability (**Figure 1D**). These results showed that bFGF remains a crucial supplement for maintaining rapid cell proliferation and robust differentiation capability. Small molecule cocktails such as ACY are desirable as a substitute for bFGF, however, there was a subsequent unexpected challenge in that cells maintained with ‘20%FBS+ACY’ exhibited stronger cell adhesion, requiring over 20 min of cell dissociation treatment after the 20th passage (data not shown). The exploration of appropriate combinations and concentrations of each inhibitor could resolve this cell dissociation problem. Further, if these compounds can be replaced with food-grade materials, they could be utilized as effective growth promoters and factors for maintaining adipogenic capability, yet also keep costs lower and thus potentially improve regulatory acceptability.

In terms of adipogenic differentiation, three different adipogenic media which were previously reported were tested in the present study (39–41). Media2 contains commonly used components for inducing adipogenesis, including insulin, IBMX (isobutylmethylxanthine), rosiglitazone, and dexamethasone (Dex) (40). Media3 includes only two inducers, insulin and rosiglitazone additions to the essential minimum required for adipogenesis (41). Media1 is based on a previously reported medium (39), to which Intralipid, has been added. Media1, which contained Intralipid, demonstrated the most efficient lipid accumulation in our isolated pDFAT cells. (**Figure 1D**). Intralipid is a fat emulsion composed of soybean oil and egg yolk phospholipids (57). It is hypothesized that egg yolk phospholipids, which contain lecithin and fatty acids, significantly enhance adipogenic differentiation (58,59). Therefore, it can be inferred that these components contribute substantially to the highest lipid accumulation among three different adipogenesis media studied in the present research. Future research should focus on the consideration of lipid-based additives that can balance the regulation of fatty acid composition with the promotion of efficient fat accumulation.

Thiamine-HCl supplementation led to the statistically significant increase of 4-methyl-5-thiazoleethanol (**Figure 5A**), and there is a tendency for thiazole, tetrahydro-, which is predictably identified, to be induced; however, no statistically significant difference was observed (**Table 1**). On the other hand, thiols and thiophenes which were supposed to be derived from thiamine, were not detected in this experimental setup. Higher sensitivity detection methods, such as GC-MS/MS, could potentially confirm additional thiamine degradation products in the cooked cultivated fat supplemented with thiamine-HCl. Furthermore, it’s possible that the amount of cell sample influences the detection of sulfide-containing VOCs. Therefore, if we prepare the cell samples through suspension culture and provide a larger amount cells, such as on a gram scale, we may be able to detect such VOCs (15).

L-methionine supplementation enhanced the production of methional, which emits a potato-like aroma (28,29) during heating/cooking. However, it was implied that L-methionine supplementation has the possibility to promote the generation of methanethiol, which is predictably identified, a degradation product of methional (60) **(Table 2)**. Methanethiol is described as having an onion-like odor, and at higher concentrations, it can be perceived as an unpleasant smell. To fully assess the impact of L-methionine on the aroma profile of cultivated pork fat, further studies incorporating descriptive sensory panels are required.

**Table 2.**
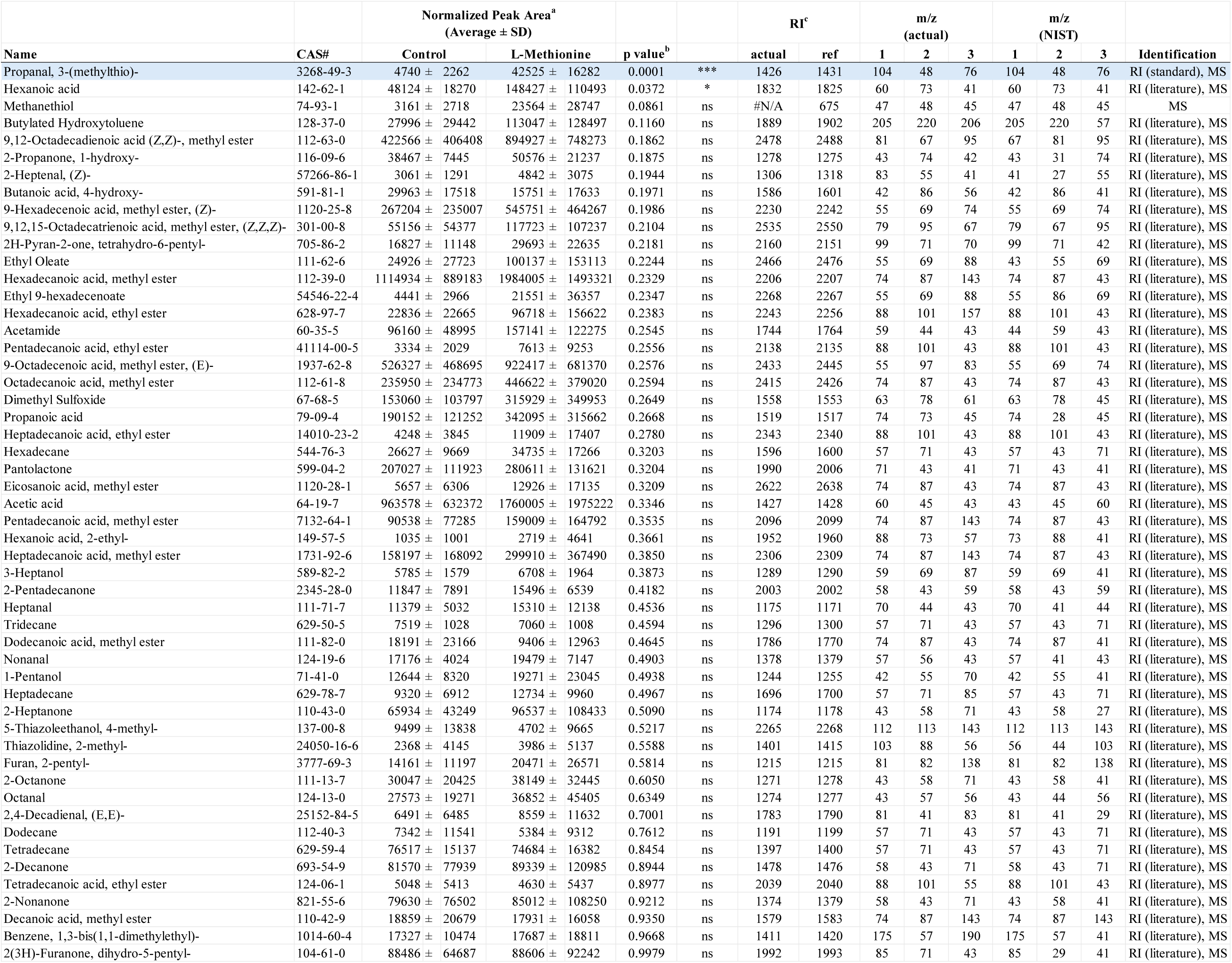
All the Volatile Organic Compounds (VOCs) found in 120℃ heated cultivated fat maintained with non-supplemented media (Control) and supplemented with 5 mM L-methionine. a) Normalized peak area was calculated by the following formula: [{(Peak area of target compound)/(Peak area of 2-Methylheptan-3-one)}*Average peak area of 2-Methylheptan-3-one]/(Peak area of Naphthalene-d8)*(Average peak area of Naphthalene-d8) and divided by the dried cell mass (mg) after baking. b) Statistical significance was determined using unpaired t-test (*, *** denote P < 0.05 and P < 0.001, respectively). c) All VOCs, except for highlighted in blue, were tentatively identified with both Mass Spec (MS) data and Retention Index (RI). Regarding the compounds shown as ‘MS, RI (standard)’ in the ‘identification’ column, RI was obtained by the authentic standard injection. RI of the other VOCs were taken from the literature which used DB-Wax column, these are considered as tentatively identified. The identification of Methanthiol is suspectable since it was identified by only MS.

**Table 3.**
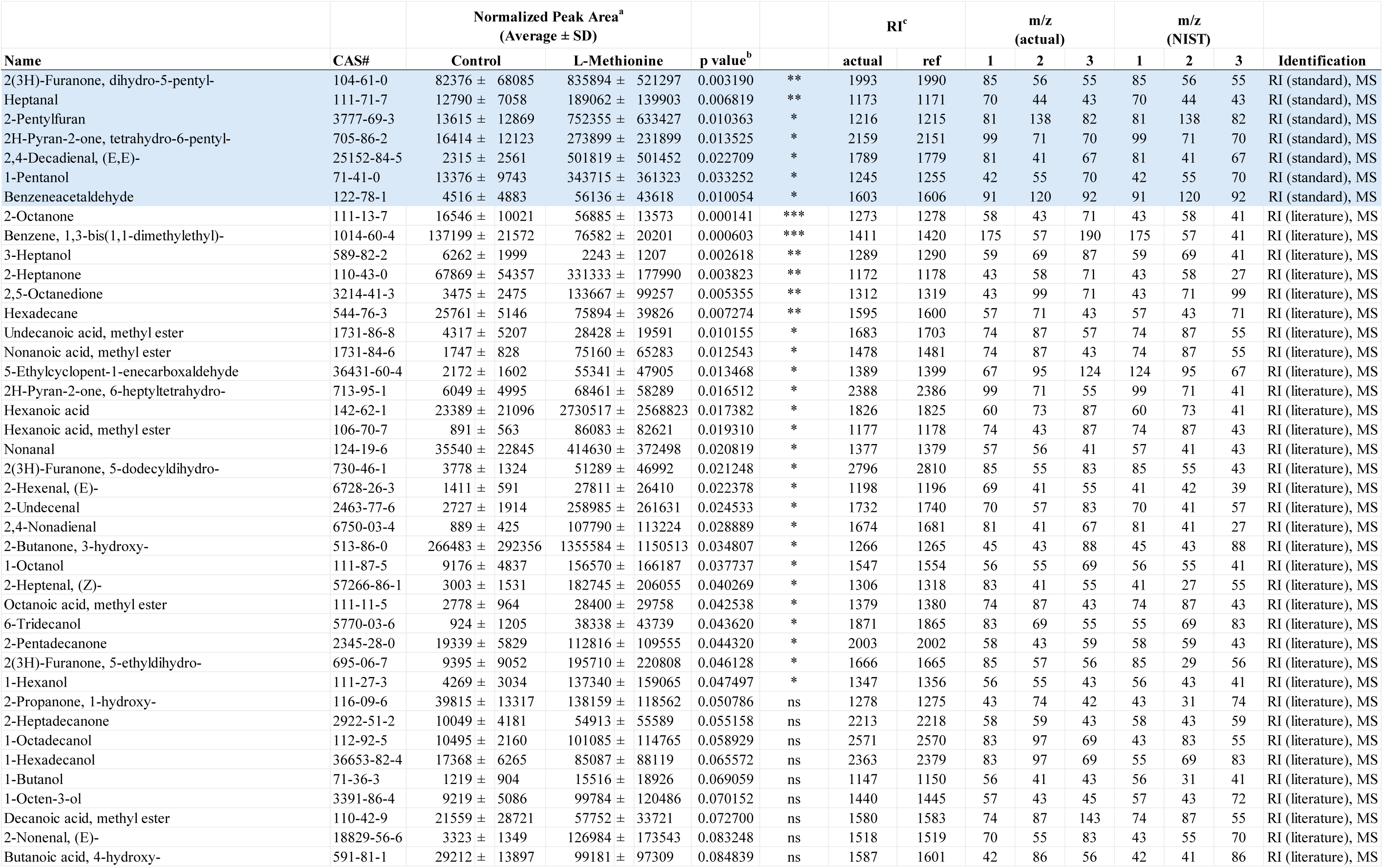
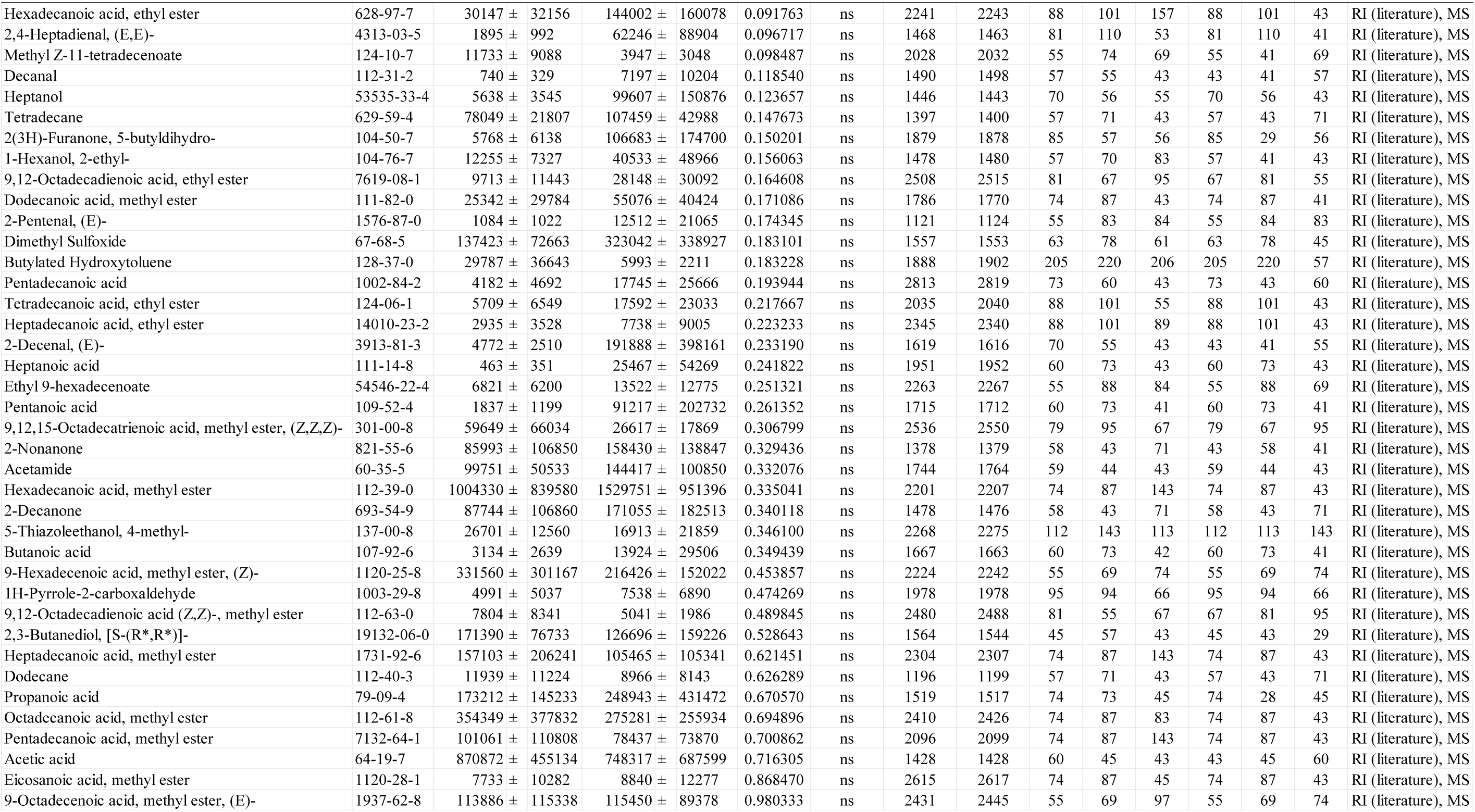
All the Volatile Organic Compounds (VOCs) found in 120℃ heated cultivated fat maintained with non-supplemented media (Control) and supplemented with 3 mg/mL myoglobin. a) Normalized peak area was calculated by the following formula: [{(Peak area of target compound)/(Peak area of 2-Methylheptan-3-one)}*Average peak area of 2-Methylheptan-3-one]/(Peak area of Naphthalene-d8)*(Average peak area of Naphthalene-d8) and divided by the dried cell mass (mg) after baking. b) Statistical significance was determined using unpaired t-test (*, **, *** denote P < 0.05, P < 0.01 and P < 0.001, respectively). c) All VOCs, except for highlighted in blue, were tentatively identified with both Mass Spec (MS) data and Retention Index (RI). Regarding the compounds shown as ‘MS, RI (standard)’ in the ‘identification’ column, RI was obtained by the authentic standard injection. RI of the other VOCs were taken from the literature which used DB-Wax column, these are considered as tentatively identified.

Myoglobin in plant-based meat increases the formation of lipid oxidation products during heating, which increases the flavor complexity and is linked to characteristics such as a serum-like taste and a metallic mouthfeel (34,61). We examined whether supplementation with myoglobin would enhance the formation of the lipid-derived aroma compounds in cultivated fat. Myoglobin significantly enhanced the formation of aldehydes, alcohols, furans, lactones and some of ketones and fatty acids (**Figure 6**). Notably, (E,E)-2,4-decadienal, a characteristic aroma compound known for its association with the deep-fat, meaty aroma of cooked meat, was reported to be 22 times greater than the non-supplemented condition (62) (**Figure 5C**). Additionally, lactones, such as γ-nonalactone and δ-decalactone, which imparts coconut-like, peachy aroma (63,64), were each statistically significantly enhanced by 7.6 times and 12 times greater than the myoglobin-supplemented conditions. Heptanal, 1-pentanol and predictably identified aldehydes such as hexanal which exhibits fatty aroma (65,66) also were enhanced. Hexanal has been associated with off-flavors that may be perceived as unpleasant at certain concentrations (65,67) (**Table 3**).

The results of the fatty acid analysis of the cultivated fat prior to cooking showed that the addition of myoglobin did not affect the fatty acid composition. Given the significant impact of myoglobin observed in this study, the combination of myoglobin and cultivated fat holds potential for altering aroma profiles (34,68). Additionally, in muscle satellite cells, it has been reported that myoglobin promotes cell proliferation (69). However, considering the cytotoxicity effects of iron in cell culture (70), the post-harvest addition of myoglobin might be a more effective depending on the cell type. Fatty acid analysis showed that cultivated fat contained 17.1% linoleic acid (**Figure 4A**). Which based on previous reports is consistent with the addition of Intralipid^35^. Decomposition of linoleic acid generates aroma compounds such as (E,E)-2,4-decadienal, 2-pentylfuran and 1-pentanol, aligning with the results observed in the present study (71–74). Thus, the fatty acid composition produced by Intralipid can be interpreted as indirectly contributing to the content of aroma compounds.

As part of future work, exploring the addition of supplements, such as L-glutamic acid or inosine monophosphate to enhance the umami taste, and linoleic acid rich edible oils provide further strategies to modify the flavor and nutritional profile of cultivated foods. For example, sunflower oil, which contains 44-75% linoleic acid (11,75,76), may enhance adipogenesis as well as modulate texture and aroma profile (77). Furthermore, investigation of masking supplements to prevent the formation of off-flavors, such as hexanal which was enhanced by myoglobin in this study, and the adjustment of myoglobin concentrations are considered crucial for further enhancing desirable odor changes. Additionally, testing various combinations and concentrations of supplements could yield other insights to provoke desirable aromas. There is a possibility that both the individual effect of ribose supplementation and its combination with other flavor precursors could synergistically enhance the formation of aroma volatiles **(Figure S2, S3)**. Furthermore, the potential to accumulate beneficial nutrients in cells, nutrients not typically found in conventional meat from livestock, is an intriguing avenue for this line of research. Such approaches may have potential not only with porcine cells but also in cells from other tissues and other species, such as bovine **(Figure S4)**, as well as in muscle cells or undifferentiated (e.g., stem) cells. The present study provides evidence that the amino acid and vitamin contents, as well as the aroma volatile profiles, of cultivated fat can be systematically altered by modulating media components, thereby enhancing the potential of cultivated fat as a food ingredient.

## 4. Material and Methods

### 4.1. Cell Isolation

Dedifferentiated porcine (*Sus domesticus*) cells (pDFAT) were isolated from the belly (subcutaneous fat) of a 93-day-old female Yorkshire pig (DOB: 10/18/2021) as previously described (36). Dedifferentiated bovine (*Bos taurus*) cells (bDFAT) were isolated from the tailhead fat tissue (subcutaneous fat) of a 604-day-old male Angus/Holstein cross steer (DOB: 09/25/2022). Briefly, fresh bovine adipose tissue was minced and digested in 0.2% collagenase (LS004176; Worthington Biochemical, Lakewood, NJ) dissolved in DMEM/F12 (11320033; Thermo Fisher, San Jose, CA), supplemented with 10% of antibiotic-antimycotic (100X) (15240062; Thermo Fisher) and 0.75% of 10% Pluronic F-68 (24040032; Thermo Fisher, San Jose, CA), for 1.5 h at 37 °C with shaking. The digest was filtered through 750 µm and then 300 µm cell strainers and centrifuged at 400 g for 5 min to collect mature adipocytes from the top layer of the supernatant. The lipid-rich layer was transferred to a tissue culture flask and incubated to allow stromal vascular cells to adhere, thereby separating them from the mature adipocytes. After 2 days, the floating lipids were transferred to a new tissue culture flask containing fresh media to initiate ceiling culture. Once dense colonies of lipid-laden cells were observed, the flask was flipped back to its normal position for routine maintenance.

### 4.2. Cell Culture

Passage 9 pDFAT cells were thawed and cultured using three different growth media formulations which developed based on previous publications (35–38), here in after ‘20%FBS’, ‘20%FBS+ACY’ and ‘15%FBS+bFGF’ with of 0.25 μg/cm^2^ laminin 511-E8 (N-892021; Iwai North America Inc., San Carlos, CA) which added to the media during cell seeding, shown in (**Table S1**). Cells were repeatedly passaged, and their doubling time compared. In ‘20%FBS+ACY’, the appropriate concentration of each inhibitor was determined by cell proliferation assay, described below (**Figure S1**). Cells were maintained by passaging at ∼70% confluency and seeding between 5,000-10,000 cells/cm^2^ or stored by freezing in growth medium with 10% dimethyl sulfoxide (DMSO) (D12345; Thermo Fisher, San Jose, CA). During passages, cells were washed with Dulbecco’s phosphate buffered saline, no calcium, no magnesium, (DPBS(-)) and dissociated with TrypLE™ Express Enzyme (1X) no phenol red (12604021; Thermo Fisher) incubating at 37°C for 10 min. The cells in growth medium were then counted using a NucleoCounter NC-200™ automated cell counter (Chemometec, Allerod, Denmark). Cell Diameter was also measured by a NucleoCounter NC-200™. Passage 23 or 24 cells were used for GC-MS analysis, samples were seeded into 150 mm dishes (430499; Corning, Tewksbury, MA). Passage 3 of bDFAT cells were maintained in ‘15%FBS+bFGF’ on laminin coated surface.

### 4.3. Cell Proliferation Assay

To determine the effect of supplementation of thiamine-HCl and L-methionine supplementation during cell proliferation, DNA amount was quantified using the CyQUANT™ Cell Proliferation Assay kit (C7026; Thermo Fisher) according to the manufacturer’s instructions. Cells were seeded in 96-well plates at 8,000 cells/cm^2^. Thiamine-HCl and L-methionine treatments were administered after the cells had adhered following seeding and cultured until the cells reach to 80% confluent. For the evaluation of A 83-01, CHIR99021 and Y-27632, cells were seeded in multiple 96-well plates at 6,000 cells/cm^2^ and cultured for 24 to 120 hours. The culture medium was replaced every other day. Every 24 hours, each plate was washed three times with DPBS(-) and frozen at - 80°C. In all experiments, the DNA amount ratio was calculated based on the condition with no chemical treatment or the condition supplemented with the maximum concentration of DMSO used for solvent, which was considered as 100%.

### 4.4. Adipogenic differentiation

After pDFAT cells reached 100% confluency in growth media and remained confluent for at least 24 h, their medium was replaced with adipogenic induction medium. The three different adipogenesis media formulations were developed based on (39–41) with replacing Chemically-defined FBS replacement changed to FBS, and 1% Penicillin/Streptomycin/Amphotericin (PSA) to 100 μg/mL Primocin (**Table S1**). For all media compositions, the cells were fed every two days until Day 8. To assess the effect of thiamine-HCl and L-methionine, using the adipogenesis Media1, supplements were applied during the adipogenesis lipid accumulation phase, days 2 to 8, with day 0 defined as the day of media transition to adipogenesis induction media. Myoglobin treatment was limited to 24 hours prior to cell harvest, due to its potential cytotoxicity and inhibition of cell proliferation (78). For each added compound, the concentration that yielded the most effective lipid accumulation was selected for use in cell culture for DHS-GC-MS analysis.

### 4.5. Lipid staining

Cultured adipocytes were stained to confirm intracellular lipid accumulation. Cells were washed twice with DPBS(-) (14190144; Thermo Fisher) to avoid cell detachment and fixed with 4% paraformaldehyde (PFA) for 20 min at room temperature (RT). After fixation, cells were first rinsed with DPBS(-), then incubated at RT for 1hr with 2 μM 4,4-difluoro-1,3,5,7,8-pentamethyl-4-bora-3a,4a-diaza-s-indacene (BODIPY 493/503, D3922; Invitrogen) diluted in DPBS(-). After BODIPY incubation, cells were rinsed three times with DPBS(-) and the cell nuclei were stained with 2 μg/mL 4’,6-diamidino-2-phenylindole (DAPI, 62247; Thermo Fisher) in DPBS(-) for 15 min at room temperature. After DAPI staining, cells were rinsed twice with DPBS(-) and stored in DPBS(-).

Imaging was performed with a fluorescent widefield microscope (KEYENCE, BZ-X700, Osaka, Japan). Using this stained plate, adipogenesis efficiency was quantified by measuring normalized BODIPY intensity by Celigo Image Cytometer (200-BFFL-5C; Nexcelom Bioscience LLC, Lawrence, MA). Specifically, the average integrated intensity of BODIPY was multiplied by the area of BODIPY, and the resulting value was normalized by dividing it by the number of nuclei stained with DAPI.

### 4.6. Fat Harvest

For GC-MS analysis, cells were prepared with Adipogenesis Accumulation Media 1 supplemented with supplementation of 500 μM of thiamine-HCl (T1270; Millipore Sigma) or 5.0 mM L-methionine (M5308; Millipore Sigma) or 25 mM ribose (R7500; Millipore Sigma) for the lipid accumulation phase, Day 2 to 8. 3 mg/mL of myoglobin (M0630; Millipore Sigma) was added only during the last 24 hours before sample collection. The spent culture media was aspirated at Day 8, then rinsed with DPBS(-) (Thermo Fisher) 3 times. Dishes were then kept vertical for 3 min to thoroughly drain DPBS(-) and any remaining media. Once excess DPBS(-) was aspirated, the adipocytes were harvested using a cell lifter (08-100-240; Fisher Scientific), then transferred into a pre-weighed 2.0 ml tube. Samples were stored at −80°C.

### 4.7. Metabolite Analysis

#### 4.7.1. Metabolite Extraction

Cells were washed in ice-cold 0.9% NaCl (S9888; Millipore Sigma) and metabolites were extracted in ice-cold 80% methanol (A456-500; Fisher Chemical) in water (W6500; Fisher Chemical) containing 500 nM isotopically labeled amino acid mix (MSK-CAA-1, Cambridge Isotope Laboratories, Tewksbury, MA) as internal standards. Samples were vortexed at 4℃ for 10 min and cleared by centrifugation for 10 min at max speed at 4℃. Supernatants were transferred to a new tube and samples were evaporated to dryness using a SpeedVac vacuum held at 6℃. The resulting cell pellet was kept on ice and used to determine protein concentrations using a Pierce™ BCA Protein Assay Kits (23225; Thermo Fisher) according to the manufacturer’s instructions.

#### 4.7.2. LC-MS Metabolite Analysis and Data Processing

Metabolite profiling was conducted on an Orbitrap Exploris 240 bench top orbitrap mass spectrometer equipped with an OptaMax™ NG Ion Source and a H-ESI probe, which was coupled to a Vanquish Horizon binary pump LC system (Thermo Fisher). External mass calibration was performed using the standard calibration mixture every 7 days and data acquisition used the Easy IC internal calibrant to maintain mass accuracy <1 ppm. To quantify methionine and the derivatives, samples were reconstituted in water such that the final concentration was equivalent to 1 μg/uL protein, and 2 μL were injected onto an Atlantis Premier BEH Z-HILIC VanGuard FIT Column: 1.7 µm, 2.1 mm × 150 mm (Waters Corporation, Milford, MA). Mobile Phase A was a 10 mM ammonium carbonate; Mobile Phase B was acetonitrile (A955-500; Fisher Chemical). The column oven and autosampler tray were held at 25℃ and 4℃, respectively. The chromatographic gradient was run at a flow rate of 0.175 mL/min as follows: 0-20 min: linear gradient from 80-20% B; 20-20.5 min: linear gradient form 20-80% B; 20.5-28 min: hold at 80% B. The mass spectrometer was operated in full-scan, polarity-switching mode, with the spray voltage set to 3.8 and –2.3 kV for positive and negative mode, respectively, the heated capillary held at 300℃, and the H-ESI probe held at 225℃. The sheath gas flow was set to 35 units, the auxiliary gas flow was set to 7 units, and the sweep gas flow was set to 1 unit. MS data acquisition was performed in a range of m/z = 70–900, with the resolution set at 120,000, the AGC target at 1×106, RF lens at 70%, and the maximum injection time at 100 msec. To quantify thiamine and thiamine pyrophosphate, samples were reconstituted to an equivalent of 0.5 μg/μL protein, and 5 uL injected onto a Luna PFP(2) LC column, 3 µm, 2 mm x 100 mm (Phenomenex). Mobile Phase A was water with 0.1% formic acid (5330020050; Thermo Fisher); Mobile Phase B was acetonitrile. The column oven was held at 30℃, and the autosampler was 4℃. The chromatographic gradient was run at a flow rate of 0.25 mL/min as follows: 0-2 min: hold at 2% B; 2-7.5 min: linear gradient from 2-60% B; 7.5-8.5 min: linear gradient from 60-100% B; 8.5-10.5 min: hold at 100% B; 10.5-15 min: hold at 2% B. The mass spec was operated in full-scan, positive ion mode with a spray voltage of 3.8 kV. All other parameters were identical to the HILIC method described above.

Relative quantitation of metabolites was performed with Skyline using a mass tolerance of 5 ppm (79). Compound ID was confirmed by referencing an in-house spectral library containing retention times built with authentic chemical standards. Data were subject to predefined quality-control parameters: CV (s.d./mean peak area across multiple injections of a representative (pooled) biological sample) below 0.25; and R (linear correlation across a three-point dilution series of the representative (pooled) biological sample) greater than 0.90. Data presented are area ratios (raw peak area endogenous metabolite/ raw peak area internal standard) normalized to total protein content.

## 5. Fatty Acid Analysis

Lipid extractions were performed using a scaled-down methyl tert-butyl ether (MTBE)-based method as described in a previous study (36). Additionally, 20 µL of a 6 mg/mL solution of nonadecanoic acid (N5252; Millipore Sigma) in hexane (139386; Millipore Sigma), measured with a glass syringe was added as an internal standard. The doubly extracted MTBE phase was then dried with nitrogen gas. The lipids were saponified by adding 3 mL of 0.5 M sodium methoxide in methanol (92446; Millipore Sigma) followed by 30 min incubation at 55°C. After cooling to room temperature, 3 mL of 14% boron trifluoride/methanol (15716; Millipore Sigma) was added, and the mixture was incubated again for 30 min at 55°C to methylate the sample. After cooling to room temperature again, the entire liquid content was transferred to a new 15 mL polypropylene centrifuge tube. Then, 2 mL of a saturated sodium chloride solution was added and vortexed, followed by the addition of 2 mL of hexane (139386; Millipore Sigma), which was vortexed again. The aqueous and organic phases were separated by centrifugation at 3,500 rpm for 5 min. The upper organic phase was transferred to a sample vial (26590; RESTEK) for gas chromatography-flame ionization detection (GC-FID) analysis.

Fatty acid composition analysis was performed using an Agilent 6890N gas chromatograph (Agilent Technologies, Santa Clara, CA) equipped with a flame ionization detector (FID) and a capillary column Select FAME (CP7430; Agilent Technologies, 100 m x 0.25 mm x 0.25 µm). The injection volume was 1.0 µL, with an inlet temperature of 250°C and a split ratio of 1:20. Helium was used as the carrier gas. The GC oven was programmed to remain at 100°C for 5 min, then increased by 10°C/min up to 220°C, where it was held for 28 min, followed by a final ramp of 10°C/min to the final temperature of 250°C, which was held for 10 min. Fatty acids were identified by comparing their retention times to those of reference standards from the Food Industry FAME Mix (35077; RESTEK, Bellefonte, PA). The standard curve was generated by serial dilution of the reference standard in hexane. Concentrations were calculated proportionally using the peak area of the internal standard, nonadecanoic acid in its methylated form. The calibration curve for methyl nonadecanoate (74208; Sigma-Aldrich) was generated by serial dilution of the reference standard in hexane, which showed a coefficient of determination (R²) of 0.996, with a limit of detection of 0.732 μg/mL (LOD), with a signal-to-noise ratio (S/N) greater than 3, which is the same with a limit of quantification (LOQ). Samples yielding peak areas within the linear range were used for data analysis. The peaks have lower than S/N = 3 were considered as not detected (n.d.).

## 6. Volatile Compound Analysis

### 6.1. Dynamic Headspace GC/MS

Volatile chemistry analysis was conducted following as previously described(13) with the exception of a modified oven temperature ramp rate, 5°C/min to 250°C. Data was acquired in scan mode ranging from 35 m/z to 300 m/z. Samples were prepared by weighing a 60 mg cell pellet into a 20 mL headspace vial (23087; RESTEK, Bellefonte, PA). As an internal standard, 1 µL of 2-Methylheptan-3-one (A284658; AmBeed, Arlington Heights, IL), prepared as 5 mg/mL in methanol, was injected into the 20 mL headspace. Additionally, prior to DHS baking, DHS tubes were loaded with Tenax® resin (11982; Millipore Sigma) and conditioned at 300°C for 120 mins under a constant flow of ultra-pure nitrogen. After conditioning, 1 µL of Naphthalene-d8 (31043; RESTEK), prepared as 10 µg/mL in dichloromethane, was injected into the Tenax® resin bed. Dichloromethane was removed by reconditioning the DHS tube at 75°C for 5 min under a constant flow of ultra-pure nitrogen. The Naphthalene-d8 was utilized to normalize the sample injection efficiency of GC/MS, and the 2-Methylheptan-3-one was utilized to normalize the DHS extraction efficiency.

### 6.2. GC-MS Data Processing

Chromatographic deconvolution was performed using PARADISe software version 6.1.7, which enabled batch processing of the full set of chromatograms. The software applies PARAFAC2 modeling within user-defined time intervals to resolve coeluting compounds. Intervals were defined to include the baseline on both sides of each peak. In cases where peaks appeared to overlap, a composite interval was created to encompass the full region as well as intervals for each visually distinct peak. Following deconvolution, the resulting mass spectra were matched against the NIST17 mass spectral libraries for “probable identification.” Only compounds with Match Quality rated as ‘Excellent’ or ‘Good’ were included. “Tentative identification” was further supported by comparison with published retention indices (RI) obtained using an identical column (DB-Wax) and chromatographic conditions. Finally, compound identities were confirmed using authentic reference standards. Deconvoluted peaks were normalized using the following formula. Deconvoluted peaks were normalized using the following formula:

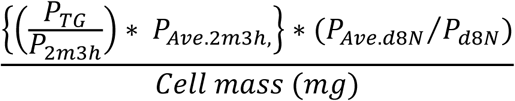

P_TG_: Peak area of target compound, P_2m3h_: Peak area of 2-Methylheptan-3-one, P_Ave.2m3h_: Average peak area of 2-Methylheptan-3-one between samples, P_Ave.d8N_: Average peak area of Naphthalene-d8 between samples, P_d8N_: Peak area of Naphthalene-d8, and Cell mass represents the baked cell mass (mg) after DHS baking.

### 6.3. Volatile Compounds Quantification

The quantification range and the detection limit of each compound were determined by the serial dilution of the following authentic standard compounds. Each compound’s quantification range, coefficient of determination of the calibration curve, limit of detection (LOD), reagent purity, and catalog number are provided in parentheses. 4-methyl-5-thiazoleethanol (7.81 to 500 µg/mL, R^2^=0.997, LOD=3.91 µg/mL, ≥98.0%, 61352), methional (1.56 to 50 µg/mL, R^2^=0.996, LOD=0.781 µg/mL, ≥97%, W274704), furan, 2-pentyl (0.078 to 50 µg/mL, R^2^=0.997, LOD=0.039 µg/mL, ≥98%, W331708), (E,E)-2,4-decadienal (0.156 to 50 µg/mL, R^2^= 0.997, LOD=0.078 µg/mL, 95%, 177510010, Thermo Scientific), 2(3H)-Furanone, dihydro-5-pentyl-(γ-Nonalactone) (0.195 to 50 µg/mL, R^2^=0.998, LOD=0.0975 µg/mL, ≥98.5%, 44542), 2H-Pyran-2-one, tetrahydro-6-pentyl-(δ-decalactone) (0.39 to 50 µg/mL, R^2^= 0.994, LOD=0.195 µg/mL, ≥98.0%, 74026), heptanal (0.195 to 50 µg/mL, R^2^=0.993, LOD=0.0975 µg/mL, ≥95%, W254002), 1-pentanol (0.0975 to 50 µg/mL, R^2^=0.999, LOD=0.0488 µg/mL, ≥99.8%, 77597), benzenacetaldehyde (0.0625 to 10 µg/mL, R^2^=0.990, LOD=0.0313 µg/mL, ≥95%, W287407), were diluted in hexane or methanol. The n-alkanes (49451-U) was used for the retention index (RI) calculations. All reagents were purchased from Millipore Sigma unless otherwise specified. In the non-supplemented control cell sample, when a peak area larger than the LOD but smaller than the limit of quantification (LOQ) was detected, the peak area was treated as the value of the LOQ, and absolute quantification was performed using the calibration curve.

## 7. Statistical Analysis

Statistical analyses were conducted GraphPad Prism 10.4.1. For comparing two groups unpaired t-tests were performed. Analysis of three or more groups was performed using one-way Analysis of variance (ANOVA) with Tukey’s post-hoc tests. Error bars and ± ranges represent standard deviations. Analysis of three or more groups in different time points or conditions was performed using two-way ANOVA with multiple comparison tests. A p-value of 0.05 was used for statistical significance. All experiments in this study were carried out with at least triplicate samples (n≥3).

## Data Availability

The DHS-GC-MS datasets generated and/or analyzed during the current study are available from the corresponding author on request.

## Ethics Declaration

Adipocyte progenitor cells were isolated from pigs and cattle in accordance with approved protocols. Porcine cells were obtained from Tufts Comparative Medicine Services (CMS), while bovine cells were sourced from Tufts Cummings School of Veterinary Medicine, with approvals under IACUC protocols #B2021-32 and #G2023-65, respectively. All procedures followed the guidelines of the United States Department of Agriculture (USDA), Office of Laboratory Animal Welfare (OLAW), Massachusetts Department of Public Health (MDPH), and the Association for Assessment and Accreditation of Laboratory Animal Care International (AAALAC). Additionally, all methods were conducted in accordance with ARRIVE 2.0 guidelines.

## Author Contributions

N.S. contributed to conceptualization, methodology, investigation, data analysis, writing – original draft, visualization. E.T.L. contributed to methodology, writing – review & editing, C.R-G. contributed to writing – review & editing, D.S.L. contributed to visualization, writing – review & editing, X.L. contributed to writing – review & editing, J.S.K.Y.Jr. contributed to cell isolation, writing – review & editing, T.L. contributed to cell isolation, writing – review & editing, A.K. contributed to data analysis, R.Y.L. contributed to investigation, Y.A.M contributed to investigation, S.C.F. contributed to methodology, investigation, writing – review & editing, D.L.K. contributed to conceptualization, manuscript drafting and editing. All the authors have read and contributed to manuscript editing.

## Acknowledgements

We gratefully acknowledge the current and past members of the Kaplan Lab and the Tufts University Center for Cellular Agriculture. We thank the Metabolomics Core at UMass Chan Medical School (RRID:SCR_027036) for assistance with experimental design, metabolite extraction, LC-MS analysis, data processing and interpretation. We thank Courtney Bogins from Tufts Comparative Medicine Services (CMS) and Eugene C. White from Cummings School of Veterinary Medicine for their valuable contributions in providing porcine and bovine adipose tissues, respectively, which were essential for cell isolation in this study. We thank Anupam Abraham for his advice on aroma optimization. We are grateful to Ajinomoto Co., Inc. for support for NS. The graphical abstract was created using BioRender. Chemical structure was designed by MolView.

## Generative AI statement

The author(s) declare that no generative AI tools were used in the preparation of this manuscript.

