## Supplementary Material for "Modulation of Nutritional Composition and Aroma Volatiles in Cultivated Pork Fat by Culture Media Supplementation"

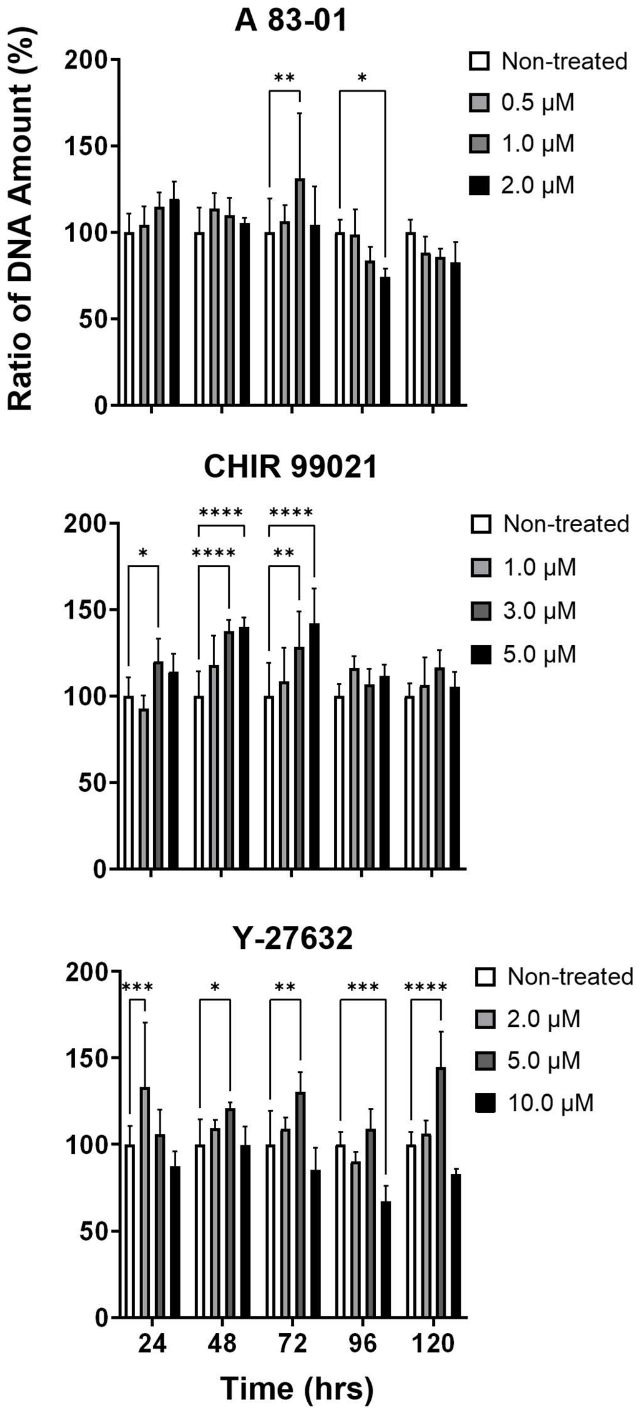


**Figure S1.** Effect of the three inhibitors, A 83-01, CHIR 99021 and Y-27632 (ACY) on cell proliferation to determine the optimal concentration for '20%FBS+ACY' proliferation media. DNA was quantified, along with the non-treated control, at each time point set to 100%. n=5 for each group. Two-way ANOVA with multiple comparisons was used to calculate significant differences. (*, **, ***, **** denote P < 0.05, P < 0.01, P < 0.001 and P < 0.0001, respectively).

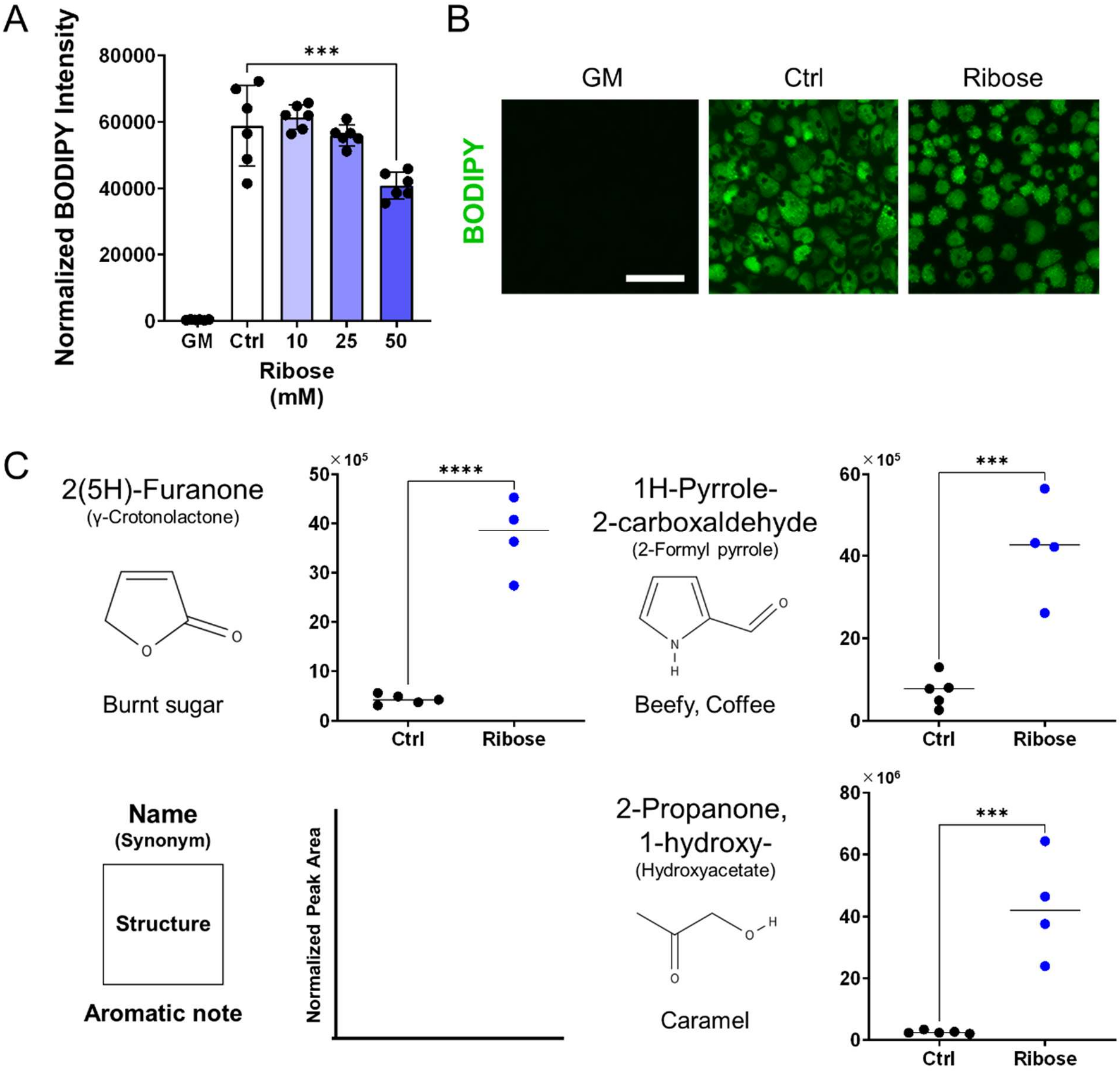


**Figure S2.** Effect of ribose supplementation during adipogenesis in pDFAT cells.
**(A)** Lipid quantification of pDFAT-derived adipocytes treated with ribose during adipogenesis lipid accumulation period. Average BODIPY integrated intensity was multiplied by BODIPY count and divided by the number of nuclei. GM; cultured in proliferation media, Ctrl; cultured in adipogenesis media without supplementation. Statistical significance was determined using one-way ANOVA followed by Tukey's test (*** denotes $P < 0.001$) compared to Ctrl. **(B)** Morphology of pDFAT-derived adipocytes cultured in adipogenesis media supplemented with 25 mM Ribose, Scale bars, 100 µm. **(C)** Major volatile organic compounds (VOCs) provoked by ribose supplementations upon cell baking. VOCs were quantified by normalizing peak areas to two internal standards and normalized to cell mass after baking. 4 replicates include biological duplicates. Statistical significance was determined using unpaired t-test (***, **** denote $P < 0.001$ and $P < 0.0001$, respectively). Compounds were identified by retention index (RI) and Mass Spec referencing NIST17.

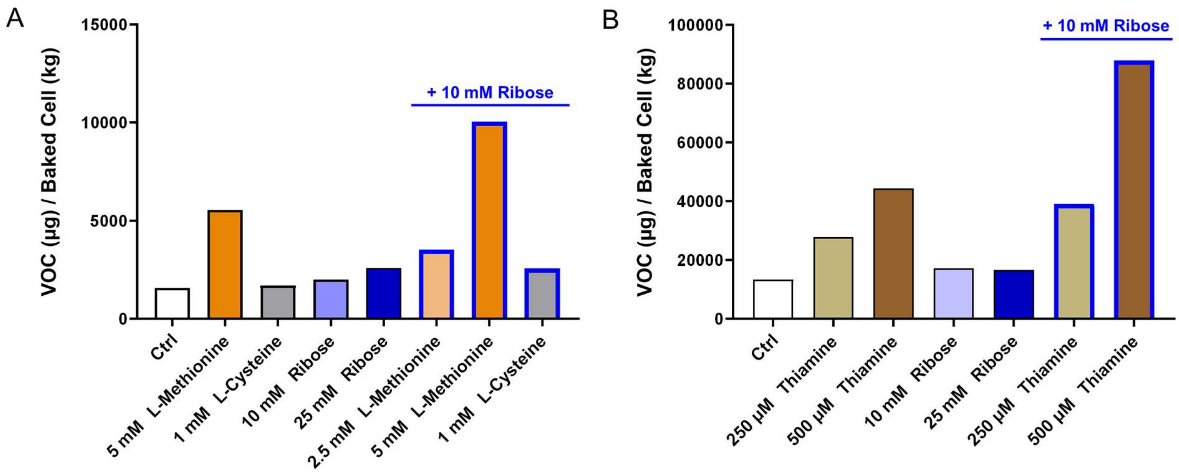


**Figure S3.** Quantification of **(A)** methional and **(B)** 4-methyl-5-thiazoleethanol under different media supplements and the combinations (n=1). Volatiles were quantified by normalizing peak areas to two internal standards and converting the values to mass using authentic standard curves. The resulting concentrations were further normalized to cell mass after baking. 'Ctrl' indicates the non-supplemented cell condition. Volatiles were identified by retention index (RI) and Mass Spec referencing NIST17 compared with authentic standards.

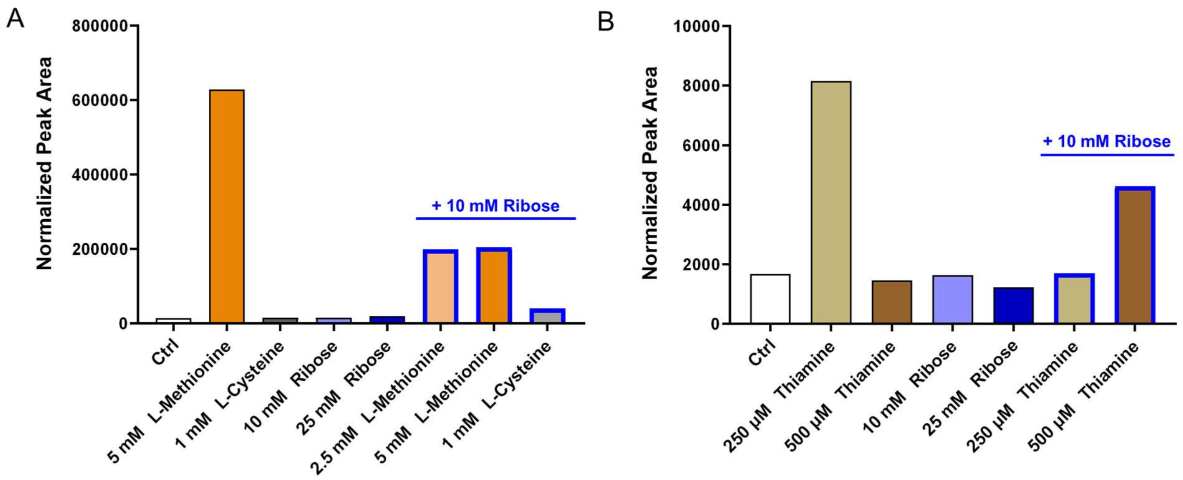


**Figure S4.** Quantification of **(A)** methional and **(B)** 4-methyl-5-thiazoleethanol upon baking bovine dedifferentiated fat cells (bDFAT) under media supplements and the combinations (n=1). VOCs were quantified by normalizing peak areas to two internal standards and normalized to cell mass after baking. Compounds were identified by retention index (RI) and Mass Spec referencing NIST17 compared with authentic standards.

**Table S1.** Cell Proliferation and Adipogenesis Media Composition

a) 3-(6-Methyl-2-pyridinyl)-N-phenyl-4-(4-quinolinyl)-1H-pyrazole-1-carbothioamide

b) 6-[[2-[[4-(2,4-Dichlorophenyl)-5-(5-methyl-1H-imidazol-2-yl)-2-pyrimidinyl]amino]ethyl]amino]-3-pyridinecarbonitrile

c) (R)-(+)-trans-4-(1-Aminoethyl)-N-(4-Pyridyl)cyclohexanecarboxamide dihydrochloride

| Component | Concentration | Supplier | Catalog No. |
|---|---|---|---|
| Proliferation Media | | | |
| 20% FBS | | | |
| DMEM-GlutaMax | N/A | Thermo Fisher | 10569-010 |
| Fetal Bovine Serum (FBS) | 20% | Thermo Fisher | A56707-01 |
| Primocin | 100 µg/mL | InvivoGen | ant-pm-1 |
| 20%FBS + ACY | | | |
| DMEM-GlutaMax | N/A | Thermo Fisher | 10569-010 |
| FBS | 20% | Thermo Fisher | A56707-01 |
| Primocin | 100 µg/mL | InvivoGen | ant-pm-1 |
| A 83-01[a] | 0.5 µM | Millipore Sigma | SML0788 |
| CHIR99021[b] | 1.0 µM | Millipore Sigma | SML1046 |
| Y-27632[c] | 2.0 µM | Millipore Sigma | Y0503 |
| 15% FBS + bFGF | | | |
| DMEM-GlutaMax | N/A | Thermo Fisher | 10569-010 |
| FBS | 15% | Thermo Fisher | A56707-01 |
| Primocin | 100 µg/mL | InvivoGen | ant-pm-1 |
| Fibroblast Growth Factor 2 (bFGF) | 10 ng/mL | PeproTech | 100-18B |

| Adipogenic Differentiation Media | | | |
|---|---|---|---|
| Media 1 | | | |
| Induction Media (first 2 days) | | | |
| DMEM-GlutaMax | N/A | Thermo Fisher | 10569-010 |
| FBS | 10% | Thermo Fisher | A56707-01 |
| Primocin | 100 µg/mL | InvivoGen | ant-pm-1 |
| Insulin | 3 µg/mL | Millipore Sigma | I0516 |
| IBMX | 0.1 mM | AdipoGen | AG-CR1-3512 |
| Dexamethasone (Dex) | 0.3 µM | Thermo Fisher | D4902 |
| Rosiglitazone | 5 µM | TCI America | R0106 |
| Biotin | 10 µM | TCI America | B0463 |
| Calcium-D-pantothenate | 5.67 uM | TCI America | P001225G |
| Lipid Accumulation Media (additional 6 days) | | | |
| DMEM-GlutaMax | N/A | Thermo Fisher | 10569-010 |
| Primocin | 100 µg/mL | InvivoGen | ant-pm-1 |
| Insulin | 3 µg/mL | Millipore Sigma | I0516 |
| Rosiglitazone | 5 µM | TCI America | R0106 |
| Biotin | 10 µM | TCI America | B0463 |
| L-ascorbic acid 2-phosphate trisodium salt | 113 µM | FUJIFILM Wako | 323-44822 |
| Intralipid | 500 µg/mL | Millipore Sigma | I141 |
| Media 2 | | | |
| Induction Media (first 3 days) | | | |
| DMEM-GlutaMax | N/A | Thermo Fisher | 10569-010 |
| FBS | 1% | Thermo Fisher | A56707-01 |
| Primocin | 100 µg/mL | InvivoGen | ant-pm-1 |
| Insulin | 10 µM | Millipore Sigma | I0516 |
| Dex | 1.0 µM | Thermo Fisher | D4902 |
| IBMX | 0.5 mM | AdipoGen | AG-CR1-3512 |
| Rosiglitazone | 5 µM | TCI America | R0106 |
| Lipid Accumulation Media (additional 6 days) | | | |
| DMEM-GlutaMax | N/A | Thermo Fisher | 10569-010 |
| FBS | 1% | Thermo Fisher | A56707-01 |
| Primocin | 100 µg/mL | InvivoGen | ant-pm-1 |
| Insulin | 58.1 µg/mL | Millipore Sigma | I0516 |
| Rosiglitazone | 5 µM | TCI America | R0106 |
| Media 3 | | | |
| DMEM-GlutaMax | N/A | Thermo Fisher | 10569-010 |
| FBS | 3% | Thermo Fisher | A56707-01 |
| Primocin | 100 µg/mL | InvivoGen | ant-pm-1 |
| Insulin | 10 µg/mL | Millipore Sigma | I0516 |
| Rosiglitazone | 5 µM | TCI America | R0106 |
